# ZBTB18 inhibits SREBP-dependent fatty acid synthesis by counteracting CTBPs and KDM1A/LSD1 activity in glioblastoma

**DOI:** 10.1101/2020.04.17.046268

**Authors:** R. Ferrarese, A. Izzo, G. Andrieux, S. Lagies, J.P. Bartmuss, A.P. Masilamani, A. Wasilenko, D. Osti, S. Faletti, R. Schulzki, Y. Shuai, E. Kling, V. Ribecco, D.H. Heiland, S.G. Tholen, M. Prinz, G. Pelicci, B. Kammerer, M. Börries, M.S. Carro

## Abstract

Enhanced fatty acid synthesis is a hallmark of tumors, including glioblastoma. SREBF1/2 regulate the expression of enzymes involved in fatty acid and cholesterol synthesis. Yet, little is known about the precise mechanism regulating SREBP gene expression in glioblastoma. Here, we show that a novel interaction between the co-activator/co-repressor CTBP and the tumor suppressor ZBTB18 regulates the expression of SREBP genes. Our study points at CTBP1/2 and LSD1 as co-activators of SREBP genes whose complex functional activity is altered by ZBTB18. ZBTB18 binding to the SREBP gene promoters is associated with reduced LSD1 demethylase activity of H3 active marks leading to increased di-methylation of lysine 4 (H3K4me2). Concomitantly, we observed increased di-methylation of lysine 9 (H3K9me2), and decrease of the active mark H3K4me3 with consequent repression of the SREBP genes. In line with our findings, lipidomic analysis shows a reduction of several phospholipid species upon ZBTB18 expression. Our results outline a new epigenetic mechanism enrolled by ZBTB18 and its cofactors to regulate fatty acid synthesis that could be targeted to treat glioblastoma patients.

## Introduction

With a median survival of 15 months and a 5-year overall survival of 5.5%, glioblastoma (GBM) is the most aggressive and deadly type of brain tumor (Ostrom et al. 2015). Over the years, GBM eluded even the most aggressive treatments (surgery followed by chemo- and radio-therapy), largely because of the invasiveness and chemo-radio-resistance of the residual tumor cells that escape resection and the subsequent treatment (Wang et al. 2016a).

GBM tumors with a mesenchymal phenotype have been described as the most aggressive, possessing features such as invasion and therapy resistance (Phillips et al. 2006; Verhaak et al. 2010; Bhat et al. 2013; Wang et al. 2017). The acquisition of mesenchymal traits in GBM is reminiscent of the mesenchymal transition in epithelial tumors and it is now considered a hallmark of aggressive tumors (Fedele et al. 2019). Epigenetic changes in the cells are inheritable, reversible covalent modifications altering gene expression without changing the DNA sequence. Histone-modifying enzymes control this process by adding or removing acetyl or methyl groups to specific position of these proteins and thus, regulating chromatin accessibility to the transcriptional machinery. These enzymes usually interact with transcription factors, co-repressor or co-activators and represent an important therapeutic opportunity in cancer (Romani et al. 2018).

The C-terminal binding proteins (CTBP1/2) can function as co-repressors through association with DNA-binding transcription factors and recruitment of chromatin regulators such as histone deacetylases 1 and 2, (HDAC1/2), lysine-specific demethylase 1 (KDM1A/LSD1), the histone methyltransferase PRC2, and the chromatin remodeling complexes NURD (Shi et al. 2003; Boxer et al. 2014; Kim et al. 2015). Although usually described as a repressor, CTBP1/2 have been also reported to act as a co-activator through the interaction with retinoic acid receptors (Bajpe et al. 2013) or through binding to the zinc finger protein RREB1 (Ray et al. 2014). CTBP1/2 have been also implicated in tumorigenesis and epithelial to mesenchymal transition (EMT) (Di et al. 2013). In GBM, CTBP1/2 have been shown to be highly expressed compared to lower grade tumors and to be associated with poorer prognosis (Wang et al. 2016b).

KDM1A/LSD1 is a histone demethylase, which acts as co-repressor or co-activator depending on the function of its protein complex. LSD1 mostly remove mono and di-methylation of histone H3 at lysine K4 (H3H4me1/2) (Shi et al. 2004). Demethylation of H3K9me2, a transcriptional repression marker of heterochromatin, has also been reported to be an LSD1 target, especially upon interaction with nuclear receptors (Metzger et al. 2005; Garcia-Bassets et al. 2007).

Sterol regulatory element-binding proteins (SREBPs) are transcription factors which control the expression of enzymes involved in fatty acids and cholesterol biosynthesis (Horton et al. 2002). SREBF1 regulates fatty acid synthesis while SREBF2 is implicated in cholesterol production (Horton et al. 2002). Fatty acid synthesis plays an important role in cancer including GBM (Bensaad et al. 2014; Lewis et al. 2015); excess of lipids and cholesterol in tumor cells are stored in lipid droplets (LD), a hallmark of cancer aggressiveness (Menendez and Lupu 2007). In GBM, constitutive activation of EGFR (EGFRvIII mutant) leads to PI3K/AKT-dependent SREBF1 regulation with consequent increase in lipogenesis and cholesterol uptake, which can be pharmacologically blocked by inhibiting the low-density lipoprotein receptor (LDLR) (Guo et al. 2011). More works have further established a role of SREBF1 as promoter of GBM growth (Cheng et al. 2015; Geng et al. 2016; Ru et al. 2016). Blocking LD formation suppresses GBM lipogenesis and growth (Geng et al. 2016); furthermore, the activation of the SREBP pathway has been recently connected to the mesenchymal shift in GBM (Schmitt et al. 2021).

Previously, we have identified ZBTB18 as tumor suppressor, which is low expressed in GBM and GBM cell lines. Others and we have previously showed that ZBTB18 functions as a transcriptional repressor of mesenchymal genes and impairs tumor formation (Carro et al. 2010; Tatard et al. 2010; Fedele et al. 2017; Xiang et al. 2021). Here, we have identified CTBP1 and CTBP2 as new ZBTB18-binding proteins. We report that CTBP and LSD1 transcriptionally activate the expression of fatty acid synthesis genes and that such activation is opposed by ZBTB18 through the inhibition of LSD1-dependent demethylase activity and the concomitant recruitment of a new repressive CTBP-LSD1-ZNF217 complex on SREBP genes promoters.

## Results

### ZBTB18 interacts with CTBP through the VLDLS motif

With the goal to get a better insight into ZBTB18 transcriptional repressive mechanisms, we used Mass Spectrometry (MS) to identify ZBTB18 co-precipitated proteins in GBM-derived brain tumor stem cell (BTSC) lines which express ZBTB18 (BTSC3082 and BTSC268) (Figure 1-figure supplement 1A-B). Among other proteins, CTBP1 and CTBP2 appeared to be potential ZBTB18 interactors (Figure 1-figure supplement 1A-B). We decided to focus on these proteins given their role as cofactors in gene expression regulation and connection to EMT and cancer. We then validated ZBTB18 interaction with endogenous CTBP1/2 in two additional BTSCs which express ZBTB18, BTSC268 and BTSC475 (Figure 1A-B). Here, however, CTBP1 and CTBP2 IP did not efficiently co-precipitated ZBTB18 probably due to the low ZBTB18 expression and suggesting that only a fraction of CTBP is involved in the interaction with ZBTB18. CTBP2 interaction with ZBTB18 was further confirmed by MS or SILAC MS upon co-immunoprecipitation with FLAG or CTBP2-directed antibodies respectively, in SNB19 cells transduced with EV or ZBTB18 (Figure 1-figure supplement 1C-D). We then validated these results in LN229 and SNB19, upon FLAG-ZBTB18 overexpression and subsequent anti-FLAG co-immunoprecipitation (Figure 1C and Figure 1-figure supplement 1E-F). ZBTB18 protein sequence analysis revealed the presence of a putative CTBP interaction motif (VLDLS) (Figure 1D) (Nibu and Levine 2001). Substitution of the LDL amino acids into SAS amino acids by site directed mutagenesis in SNB19 cells completely abolished CTBP2 interaction with ZBTB18 (Figure 1E-F), further validating ZBTB18-CTBP interaction. Consistent with our previous study (Fedele et al. 2017), ZBTB18 affected cell proliferation, apoptosis and migration (Figure 1-figure supplement 2A-F). ZBTB18 LDL mutant (ZBTB18-mut) also appeared to have a mild effect on apoptosis and proliferation while migration was not impaired (Figure 1-figure supplement 2A-F). Expression analysis of previously validated ZBTB18 mesenchymal targets, upon overexpression of the ZBTB18-mut, showed that ZBTB18 interaction with CTBP is required for ZBTB18-mediated repression of a subset of targets (CD97, LGALS1 and S100A6, Figure 1G), while repression of ID1, SERPINE1 and TNFAIP6 type of genes appears to be independent from CTBP2 binding. Overall, these data identify CTBP1/2 as new ZBTB18 interactors in GBM cells and suggest that ZBTB18 might employ both a CTBP-dependent and a CTBP-independent mechanism to repress target genes and mediate its tumor suppressor functions.

**Figure 1.**
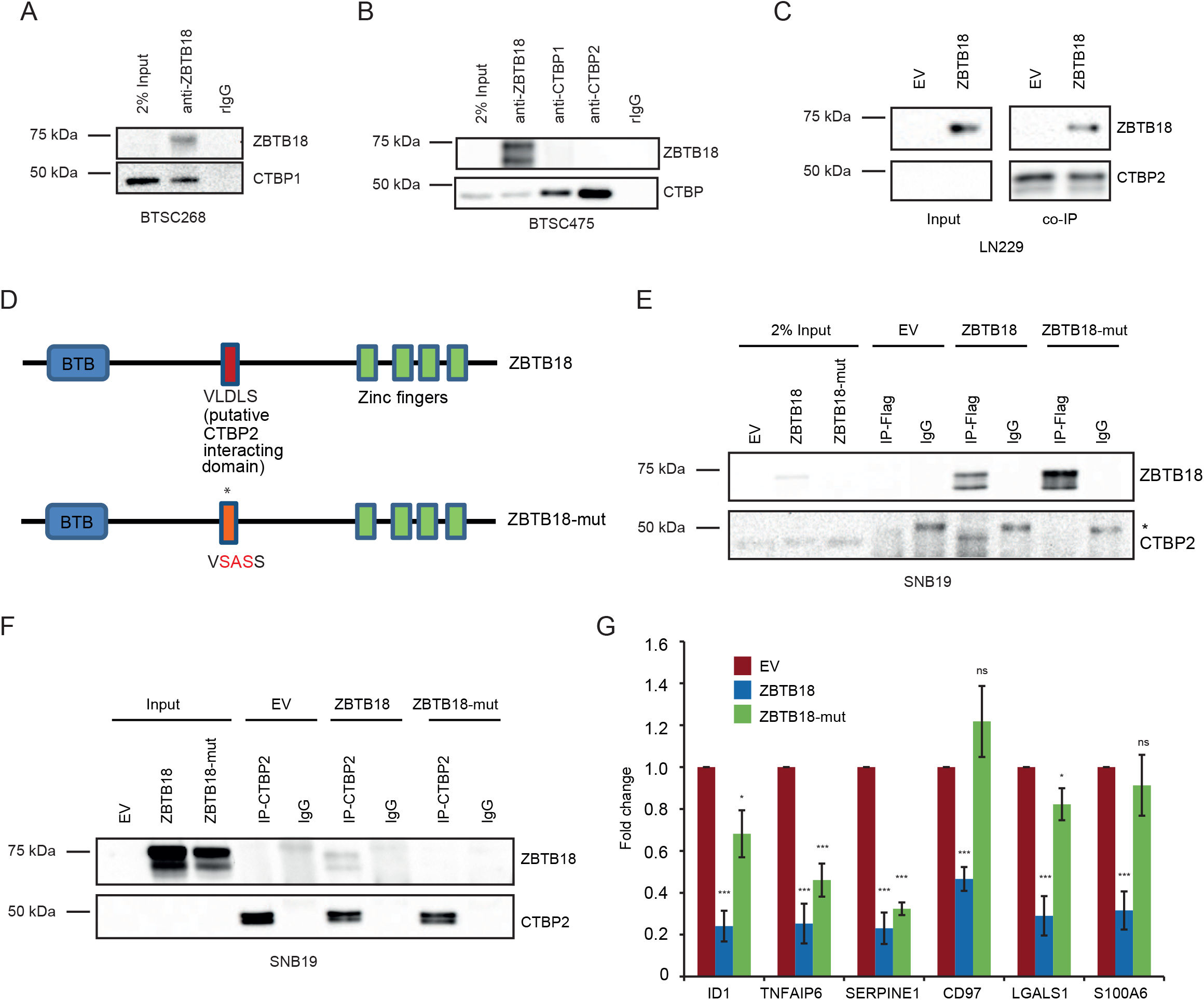
ZBTB18 interacts with CTBP through the VLDLS domain in glioblastoma cells. (**A-B**) WB analysis of endogenous ZBTB18 co-IP in BTSC268 (**A**) and BTSC475 (**B**) glioblastoma cells. (**C**) WB analysis of FLAG co-IP in LN229 cells transduced with empty vector (EV) or FLAG-ZBTB18. (**D**) Schematic representation of the ZBTB18 protein with BTB and Zinc fingers domain. The putative CTBP2 interacting motif (VLDLS) and the corresponding mutation are marked. (**E-F**) WB analysis of FLAG co-IP (**E**) or CTBP2 co-IP (**F**) in SNB19 cells expressing either ZBTB18 or ZBTB18-mut. (**G**) qRT-PCR showing expression of ZBTB18 targets upon ZBTB18 and ZBTB18-mut overexpression in SNB19 cells. n=3 biological replicates; error bars ± s.d. *p < 0.05, **p < 0.01, ***p < 0.001 by Student’s t-test. Gene expression was normalized to 18sRNA.

### ZBTB18 and CTBP2 play an opposite role in gene regulation

Since CTBP1/2 are mostly known as co-repressor, we hypothesized that CTBP1/2 could be required for ZBTB18 function. To verify this possibility, we overexpressed FLAG-ZBTB18 and knocked down CTBP2, which interacts with ZBTB18, in SNB19 cells (Figure 2A). Gene expression profiling followed by gene set enrichment analysis (GSEA) showed that both ZBTB18 overexpression and CTBP2 silencing similarly affected the expression of EMT gene signatures, suggesting an opposite role of CTBP2 and ZBTB18 in transcriptional regulation (Figure 2-figure supplement 1A). This is consistent with our reported role of ZBTB18 as inhibitor of mesenchymal signatures in GBM and with previous studies indicating that CTBP2 promotes tumorigenesis and EMT.

**Figure 2.**
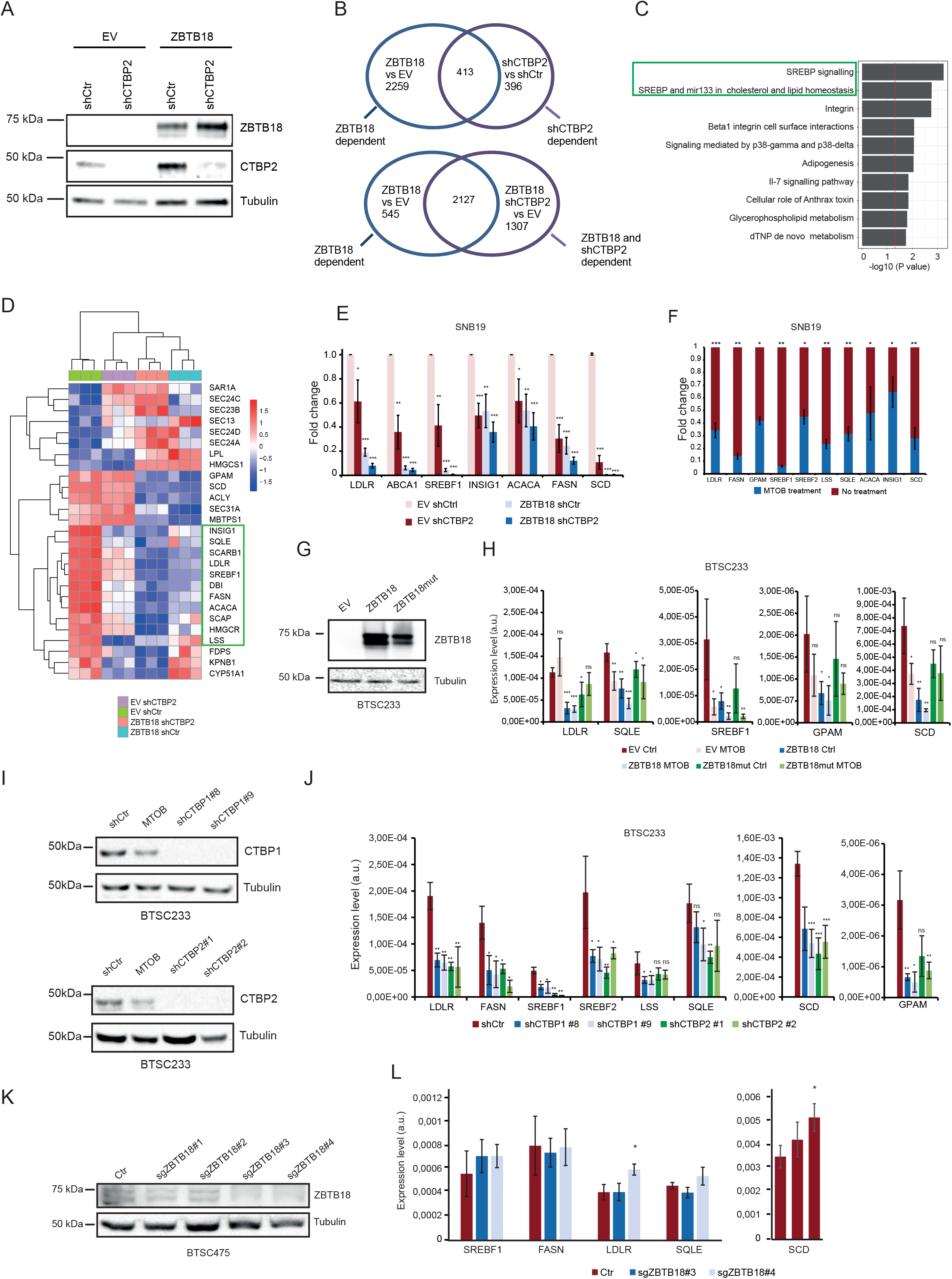
ZBTB18 and CTBP regulate the expression of SREBP genes involved in fatty acid synthesis. (**A**) WB analysis of FLAG-ZBTB18 and CTBP2 expression in SNB19 cells transduced with either FLAG-ZBTB18 or shCTBP2. (**B**) Venn analysis showing the overlap between regulated genes in ZBTB18 vs. empty vector (EV) and genes regulated in shCTBP2 vs. shCtr (upper part) or ZBTB18 shCTBP2 vs. EV (lower part). Genes were selected based on adjusted p value < 0.05 and absolute fold change > 0.5. (**C**) Top 10 downregulated consensus pathways in the overlap between shCTBP2 vs. shCtr and ZBTB18 vs. EV regulated genes (ZBTB18 vs shCTBP2) from Fisher’s exact test comparing the DEGs (adj. p value < 0.05, FC < −0.5) to the whole set of quantified genes. Processes related to SREBP signaling are highlighted. (**D**) Row-wise z-score heatmap showing the expression of SREBP signaling genes in each triplicate across the 4 conditions. Rows and columns hierarchical clustering are both based on Euclidean distance and complete clustering method was used. (**E**) qPCR validation of selected SREBP genes expression upon ZBTB18 expression and/or CTBP2 silencing in SNB19 cells. n=3 biological replicates; error bars ± s.d. *p < 0.05, **p < 0.01, ***p < 0.001 by Student’s t-test. Gene expression was normalized to 18s RNA. (**F**) qPCR analysis of selected SREBP genes in SNB19 cells treated with the CTBP2 inhibitor MTOB. n=3 biological replicates; error bars ± s.d. *p < 0.05, **p < 0.01, ***p < 0.001 by Student’s t-test. Gene expression was normalized to 18s RNA. (**G**) WB analysis of ZBTB18 expression in BTSC233 cells transduced with empty vector (EV), ZBTB18 or ZBTB18-mut, using a ZBTB18 antibody. (**H**) qPCR analysis of selected SREBP genes upon ZBTB18 overexpression and MTOB treatment. Results are presented as the mean of n=3 biological; error bars ± s.d. *p < 0.05, **p < 0.01, ***p < 0.001 by Student’s t-test. Gene expression was normalized to 18s RNA. (**I**) Western blot analysis of CTBP1 (top panel) and CTBP2 (bottom panel) expression upon silencing in BTSC233 cells. (**J**) q-RT PCR analysis of SREBP targets in BTSC233 transduced with shCTBP1 or shCTBP2 expressing lentivirus. Two independent shRNA to knock down CTBP1 or CTBP2 were used. Results are presented as the mean of n=3 biological; error bars ± s.d. *p < 0.05, **p < 0.01, ***p < 0.001 by Student’s t-test. (**K**) WB analysis of ZBTB18 expression in BTSC475 cells upon transduction with the indicated ZBTB18 sgRNAs. (**L**) q-RT PCR analysis of selected SREBP gene expression, upon ZBTB18 knockdown (sgZBTB18#3 and sgZBTB18#4). Error bars ± s.d.. Gene expression was normalized to 18s RNA.

Venn analysis of shCTBP2 and ZBTB18 regulated genes showed that about half of the genes regulated by shCTBP2 are also repressed upon ZBTB18 overexpression (Figure 2B, top panel). CTBP2 knockdown partially affected the portion of ZBTB18 regulated genes supporting the observation that ZBTB18 may regulate gene expression both in a CTBP2-dependent and independent manner (Figure 2B, bottom panel). GSEA revealed a strong loss of SREBP signaling genes expression, both upon ZBTB18 overexpression and CTBP2 silencing (Figure 2C-D and Figure 2-figure supplement 1B), according to a possible role of CTBP2 as co-activator, and of ZBTB18 as repressor. SREBP genes are involved in fatty acid synthesis. We focused on the SREBP signaling pathway given its importance in glioblastoma tumorigenesis and previous connection to EMT (Guo et al. 2011; Cheng et al. 2015; Wang et al. 2015; Yang et al. 2015; Geng et al. 2016; Ru et al. 2016; Zhang et al. 2019). Interestingly, SREBF1 activation has been recently shown to promote a mesenchymal shift in GBM (Schmitt et al. 2021). Consistent with the literature reports, GlioVis analysis (Bowman et al. 2017) showed that SREBF1 is more expressed in the most aggressive GBM subtypes (mesenchymal and classical) and correlates with patient survival (Figure 2-figure supplement 1C-D). Furthermore, we observed a strong positive correlation between CTBP2 expression in gliomas and the levels of all the SREBP genes tested (GlioVis platform, dataset CGGA (Zhao et al. 2017), Figure 2-figure supplement 2), which is in line with a possible role of CTBP2 as co-activator of SREBP genes. Validation by qPCR in SNB19 and various BTSC lines confirmed the reduction in the expression of all the genes tested upon ZBTB18 expression (Figure 2E, 2G-H and Figure 2-figure supplement 3A-B) or CTBP2 silencing (Figure 2E). In agreement with our microarray results, CTBP2 knockdown did not inhibit ZBTB18-mediated repression (Figure 2D-E) but rather it further enhanced downregulation of ZBTB18 target genes. Remarkably, treatment with the CTBPs inhibitor MTOB (Achouri et al. 2007) also resulted in a strong downregulation of SREBP genes expression in SNB19 cells (Figure 2F), while in BTSC233 it mostly affected SREBF1, SCD and SQLE expression (Figure 2H). Knockdown of CTBP1 and CTBP2, either alone or in combination, in BTSCs further proved CTBPs activating role (Figure 2I-J and Supplementary Fig.S6C-E). When MTOB treatment was combined with ZBTB18 or ZBTB18-mut expression in BTSC233, no rescue of ZBTB18 repression was observed (Figure 2H); however, ZBTB18-mut seemed to be less effective in repressing SREBP genes (Figure 2H), suggesting that MTOB might specifically affect the transcriptional activating role of CTBP without impairing ZBTB18 repressive function. Consistent with this idea, MTOB was shown to cause displacement of CTBP from its target gene promoters with no impairment of its interaction with transcription factors (Di et al. 2013). We then attempted to knock down ZBTB18 in the primary GBM cell line BTSC475 by CRISPR/Cas9. Despite the basal expression level of ZBTB18 in GBM cells, knock out with the *ZBTB18*-directed sgRNA#3 and #4 led to a detectable increase of SCD and LDLR, consistent with ZBTB18 repressive role (Figure 2 K-L).

### ZBTB18 affects lipid synthesis and reduces lipid storage

Then, we investigated whether ZBTB18 overexpression caused phenotypic changes associated with the reduction of fatty acid synthesis, due to the deregulated SREBP signaling gene expression. SNB19 cells were transduced with ZBTB18 or ZBTB18-mut and profiling of lipid species expression was performed by a targeted LC-MS method (Figure 3A). Hierarchical clustering based on lipid species relative abundance highlighted that ZBTB18-expressing cells segregated together with each other and separately from the other samples (empty vector (EV) and ZBTB18-mut) regardless of the growing conditions (normal medium or lipid-depleted medium) (Figure 3B-C). Within this cluster, significantly regulated lipids were largely underrepresented in the ZBTB18-expressing cells compared to the EV control cells; notably, the lipid starvation exacerbated this trend. Of note, among the lipids downregulated upon ZBTB18 ectopic expression, several contained unsaturated carbon chains, suggesting that the desaturation step in the biosynthesis of fatty acids was particularly affected by ZBTB18.

**Figure 3.**
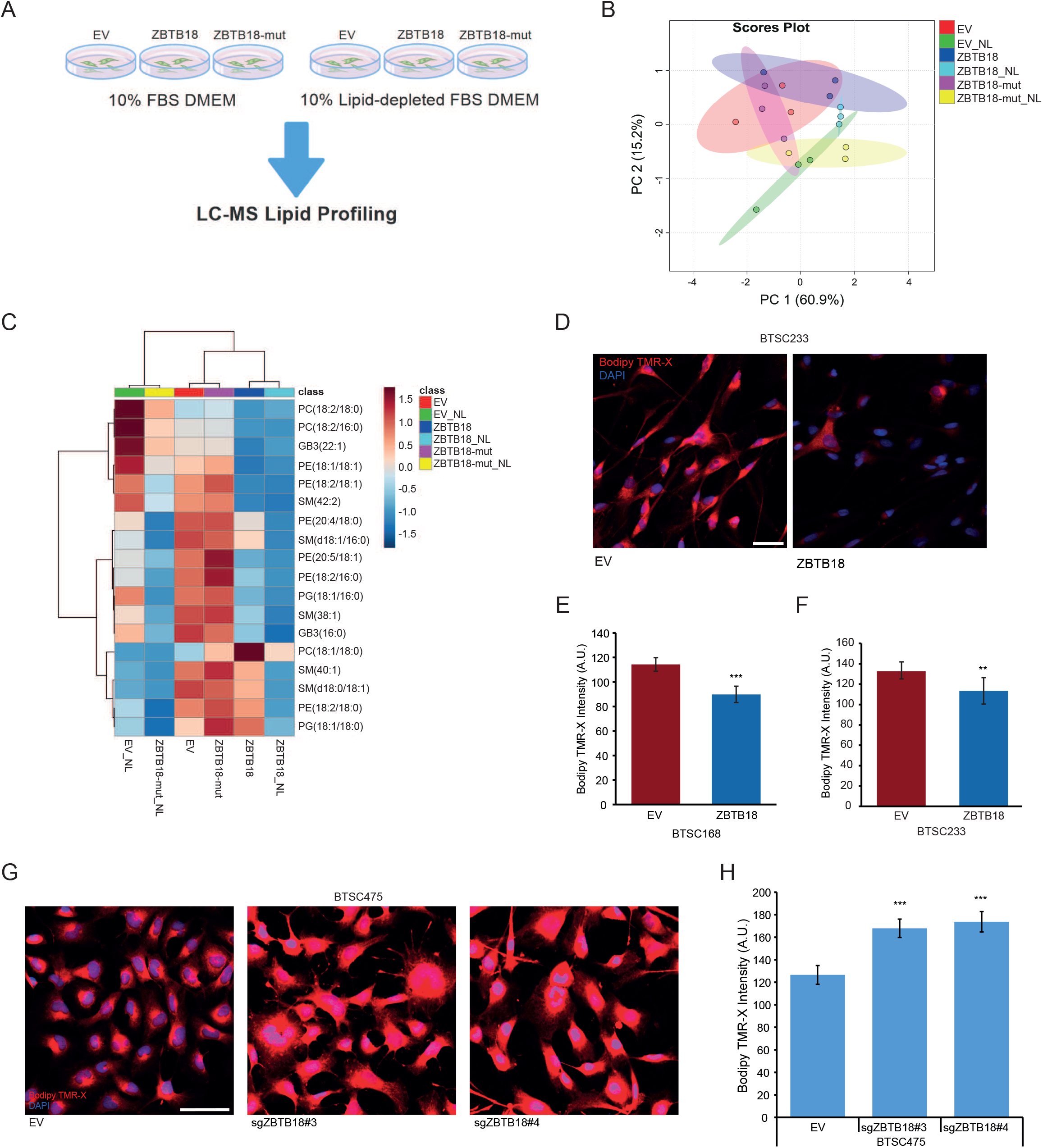
ZBTB18 overexpression alters lipid content depending on its CTBP binding capability. (**A**) Experimental flow chart of lipidomics analysis. (**B**) Principal component analysis of identified lipids in SNB19 cells transduced with empty vector (EV), ZBTB18 overexpressing vector (ZBTB18) or ZBTB18 mutant (ZBTB18-mut) and grown in the presence (no suffix) or absence of lipids (NL). n=3. (**C**) Heatmap of significantly altered (q < 0.05) lipids in SNB19 cells expressing EV, ZBTB18 or ZBTB18-mut, grown in the presence (no suffix) or absence of lipids (NL). Range-scaled z-scores are displayed. (**D**) Bodipy TMR-X lipid staining of BTSC233 cells expressing EV or ZBTB18. Nuclei were counterstained with DAPI. Scale bar: 100μm. (**E-F**) Quantification of Bodipy TMR-X lipid staining in BTSC168 (**D**) and BTSC233 (**F**). n=4 biological replicates; error bars ± s.d. *p < 0.05, **p < 0.01, ***p < 0.001. (**G**) Bodipy TMR-X lipid staining of BTSC475 cells upon ZBTB18 KO (sgZBTB18#3 and sgZBTB18#4). Nuclei were counterstained with DAPI. Scale bar: 100μm. (**H**) Quantification of Bodipy TMR-X lipid staining shown in (**G**).

To further confirm a role of ZBTB18 in lipid turnover in GBM cells, we analyzed lipid droplets content upon ZBTB18 overexpression. In both the tested primary GBM stem cell lines (BTSC168 and BTSC233), the presence of ZBTB18 led to a significant reduction of the amount of lipid droplets within the cells (Figure 3D-F). A similar effect was observed in SNB19 cells, in which the reduced number of lipid droplets upon ZBTB18 overexpression, became more evident after a 48-hour lipid starvation (Figure 3-figure supplement 1A-B). Moreover, when lipid-starved, ZBTB18-expressing cells were incubated with lipid-containing medium again, they showed a significant increment of lipid droplets albeit not fully recovering to the level of the EV controls (Figure 3-figure supplement 1A-B). This observation suggests that the loss of lipid droplets in cells expressing ZBTB18 is mostly due to the blockade of the lipid biosynthesis on which tumor cells especially rely when there are no lipid sources available in the environment. In agreement with this hypothesis, no significant difference in the uptake of fluorescently labelled palmitic acid (Bodipy-C16) was observed between ZBTB18-expressing cells and the respective controls (Figure 3-figure supplement 1C). Then, we measured lipid droplet content upon ZBTB18 knockdown; remarkably, the number of lipid droplets strongly increased upon ZBTB18 deletion with both sgRNA (Figure 3G-H).

Taken together, these data suggest that ZBTB18 plays an important role in controlling lipid metabolism of GBM cells.

### ZBTB18 and CTBP2 map to SREBP promoter regions

To characterize the dynamics of CTBP2 and ZBTB18 on gene targets regulation, we mapped the genome wide distribution of CTBP2 and ZBTB18 by chromatin immunoprecipitation coupled to deep sequencing (ChIPseq) in SNB19 cells. CTBP2 ChIP was performed with a CTBP2 antibody to precipitate the endogenous CTBP2 in absence or in presence of ZBTB18 (EV_CTBP2 and ZBTB18_CTBP2) while ectopic ZBTB18 was immunoprecipitated with FLAG antibody (EV_FLAG and ZBTB18_FLAG) (Figure 4A).

**Figure 4.**
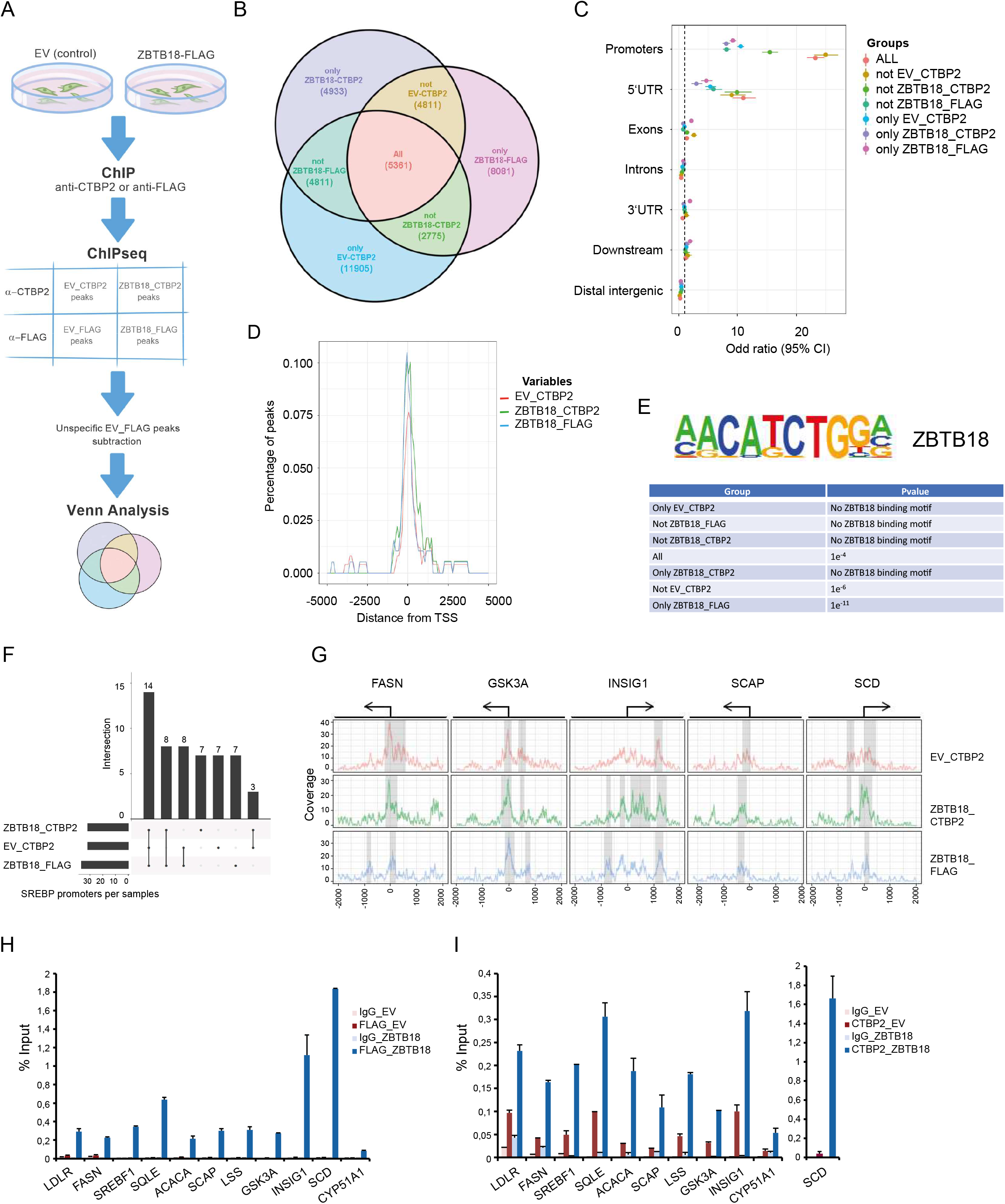
CTBP2 and ZBTB18 map to SREBP gene promoters. (**A**) Experimental flow chart of the ChIP-seq analysis. (**B**) Venn diagram showing the overlap between the 3 consensus peak sets from EV_CTBP2, ZBTB18_CTBP2 and ZBTB18_FLAG. The percentage of overlap is indicated in red. (**C**) Enrichment of peaks on promoter regions depicted on a forest plot from Fisher’s test analysis. Dots indicate the Odd ratio and lines indicate the 95% confidence interval. (**D**) Meta-gene peak density profile showing an over-representation of peaks around SREBP genes TSS regions. Y-axis indicates the percentage of peaks in each group that overlap the genomic region depicted on the X-axis. (**E**) Homer analysis showing enrichment of the ZNF238/ZBTB18 DNA binding motif in the Venn diagram groups. (**F**) UpSet plot showing the overlap of SREBP genes having at least one peak in one of the 3 conditions: EV_CTBP2, ZBTB18_CTBP2 and ZBTB18_FLAG. (**G**) Read coverage around biological relevant selected genes, i.e. FASN, GSK3A, INSIG1, SCAP and SCD. Identified peaks are highlighted under the grey area and genes TSS are drawn on top of the plot. (**H-I**) FLAG-ZBTB18 (**H**) and CTBP2 (**I**) enrichment at the promoter of the indicated SREBP genes in SNB19 cells transduced with empty vector (EV) of FLAG-ZBTB18. ChIP was performed using control beads alone (IgG), anti-FLAG antibodies or anti-CTBP2 antibody. Graphs show representative qPCR results (n=3 technical replicates) of at least two biological replicates and are expressed in % input as indicated.

After subtracting FLAG unspecific peaks detected in SNB19-EV_FLAG, we performed Venn analysis of all the annotated peaks. We found 5361 peaks shared by all conditions suggesting that CTBP2 and ZBTB18 bind to the same genomic regions (Figure 4B). Remarkably, peaks which are in common with all the conditions (All), and those that are shared between ZBTB18 and CTBP2 ChIPs when ZBTB18 is expressed (Not EV_CTBP2) were strongly enriched at promoter regions in close proximity to the transcription start (Figure 4C-D). This suggests that, while CTBP2 and ZBTB18 share common sites (All), a fraction of CTBP2 can be recruited to new promoter regions when ZBTB18 is expressed (Not EV_CTBP2). *In silico* analysis of consensus binding motif by HOMER showed enrichment for ZBTB18 in both groups (Not EV CTBP2 and All), in addition to the “only ZBTB18_FLAG” group (Figure 4E). Promoter peaks belonging to the group “Not ZBTB18_CTBP2”, which are not shared by ZBTB18 and CTBP2 upon ZBTB18 expression, do not contain the ZBTB18 motif, suggesting that these peaks could be unspecific (Figure 4E). Analysis of published ChIP-seq datasets using ReMap (Cheneby et al. 2019) showed a good overlap between shared CTBP2 and ZBTB18 promoter peaks, with peaks for known CTBP2 interactors such as NCOR1, ZNF217 and LSD1/KDM1A (p=0, Figure 4-figure supplement 1A and C). When focusing on SREBP gene promoters, we observed a strong consensus between peaks identified in ZBTB18 (FLAG) and CTBP2 ChIPs (Figure 4F and Supplementary Table S1), again suggesting that CTBP2 and ZBTB18 play a direct role in SREBP genes transcription. Furthermore, SREBP gene peaks common to ZBTB18 and CTBP2 ChIPs, matched with genes regulated by CTBP2 silencing and ZBTB18 overexpression in our gene expression analysis (INSIG1, SREBF1, FASN, ACACA, LDLR, DBI, SCAP, SQLE and SCD; Figure 2D-E and Figure 4G and Supplementary Table 1). Furthermore, ReMap analysis revealed a good consensus between regions shared by ZBTB18 and CTBP2 and SREBP peaks (p=0, Figure 4-figure supplement 1B-C). Together, these results further reinforce the notion that CTBP2 and ZBTB18 bind to a common gene promoters which include SREBP genes. In addition, this constitutes the first genome wide mapping of CTBP2 and ZBTB18 in GBM cells.

We further confirmed the binding of ZBTB18 to the promoter of a set of SREBP genes by qChIP analysis glioblastoma cells. Upon overexpression in SNB19 cells, ZBTB18 is recruited to the promoter region of all analyzed genes but not of CYP51A1, which is a preferential target of CTBP2 alone (Figure 4H). CTBP2 is present at the promoter of SREBP genes in the absence of ZBTB18 (Figure 4I) where according to our microarray analysis and qPCR results it is responsible for their transcriptional activation. Interestingly, we observed a consistent increase of CTBP2 levels at the promoter of SREBP genes in the presence of ZBTB18, but not when ZBTB18-mut was expressed, in both BTSC168 and SNB19 cells (Figure 4-figure supplement B and D), suggesting that the interaction between ZBTB18 and CTBP2 adds complexity to the regulatory dynamics of the latter. In line with the proposed role of ZBTB18 as a repressor of SREBP genes expression, we observed that the levels of H3K4me3, a well-known marker of transcriptional activation, decreased at the promoters of all the analyzed genes (Figure 4-figure supplement 2E).

### ZBTB18 facilitates CTBP2 binding to ZNF217, LSD1 and NCOR1

To better understand whether the presence of ZBTB18 affects CTBP2 interaction with other proteins, we performed SILAC-based MS in SNB19 cells transduced with either control vector (low-weight medium, L) or ZBTB18 (high-weight medium, H) followed by CTBP2 co-immunoprecipitation (Figure 5A). Interestingly, ZNF217, RCOR1, LSD1/KDM1A and HDAC1/2, which are well-characterized CTBP1/2 interactors, more efficiently co-precipitated with CTBP2 when ZBTB18 was expressed (H/L=1.83, 1.73, 1.6 and 1.48 respectively) (Figure 5B). CTBP2 co-immunoprecipitation in BTSC168 and SNB19 cells followed by western-blot analysis confirmed the increased binding of ZNF217, LSD1 and CTBP to each other, upon ZBTB18 expression (Figure 5C-D and Figure 5-figure supplement 1A-B). Of note, LSD1 protein levels appeared to be higher upon ZBTB18 expression (Figure 5C). Overall, these pieces of evidence suggest that CTBP2 complex dynamics and functions could change upon ZBTB18 overexpression. To verify whether ZBTB18 binding to CTBP2 is required for the recruitment or stabilization of the CTBP2 complex at SREBP gene promoters, ChIP experiments on selected SREBP genes were performed in SNB19 cells upon overexpression of ZBTB18, using LSD1 and ZNF217 directed antibodies. Here, both LSD1 and ZNF217 appeared to be more enriched at SREBP gene promoters when ZBTB18 was expressed (Figure 5E-F). Interestingly, this effect was lost or much reduced when ZBTB18mut was expressed, in both SNB19 and BTSC168 cells (Figure 5G and Figure 5-figure supplement 1C-D). Interestingly, ChIP analysis of H3K4me2 and H3K9me2 binding at selected SREBP gene promoters, upon ZBTB18 or ZBTB18-mut overexpression in BTSC168, displayed an enrichment for both histone marks in ZBTB18 expressing cells (Figure 5H-I). Similarly, measurement of H3K4me2 methylation using a commercial assay confirmed that ZBTB18 but not ZBTB18-mut impaired LSD1 activity (Figure 5-figure supplement 1E). Together these data suggest that, through binding to CTBP, ZBTB18 favors the recruitment of LSD1 to the SREBP gene promoters. At the same time, LSD1 demethylase activity appears to be impaired.

**Figure 5.**
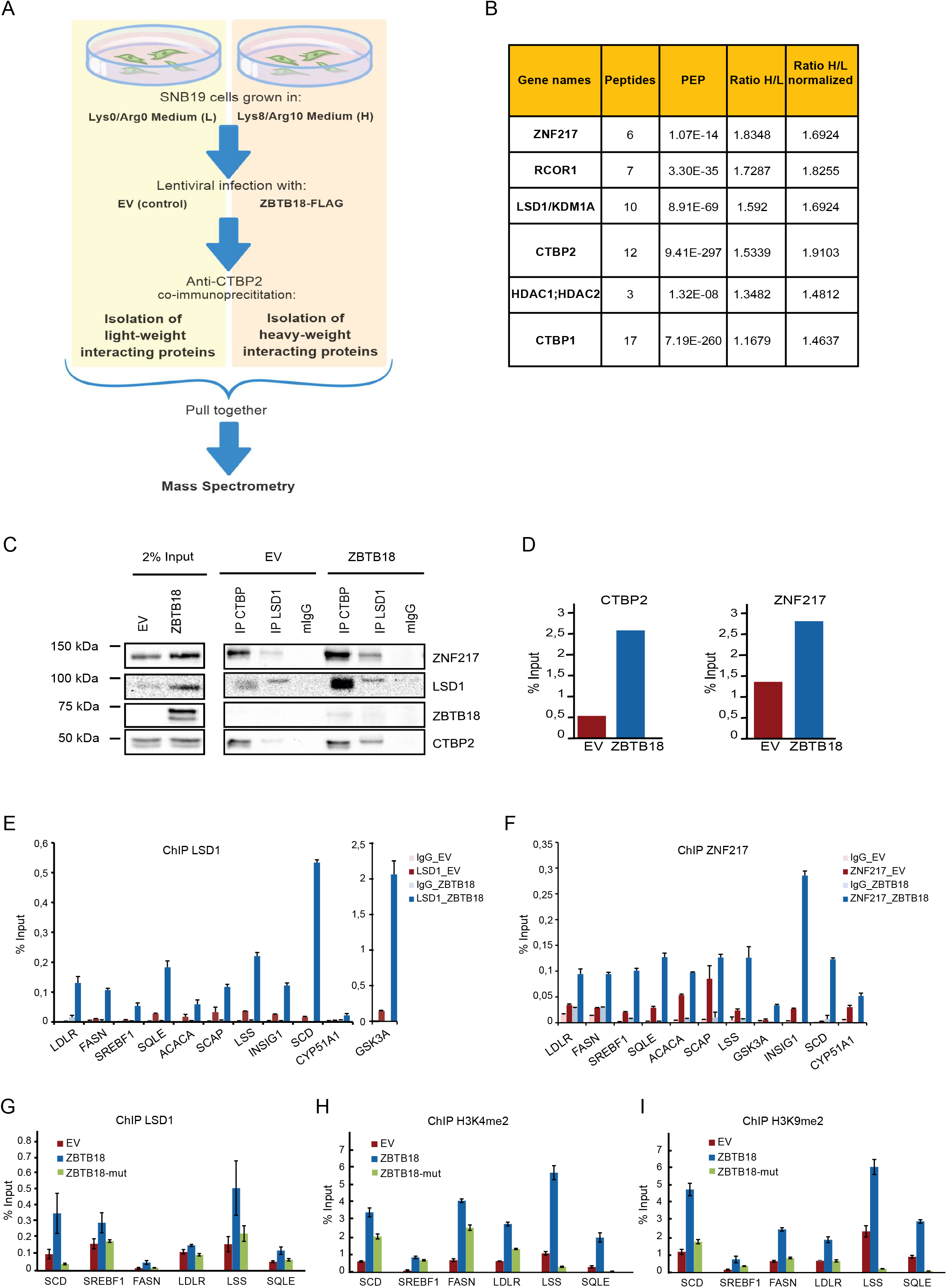
ZBTB18 recruits CTBP2 complex to the promoter of SREBP genes. (**A**) Experimental flow chart of the SILAC-based MS analysis. (**B**) List of known CTBP2 interacting proteins identified by SILAC. Binding to CTBP2 increases in the presence of ZBTB18 (H) respect to EV (L). (**C**) WB analysis of LSD1 or CTBP co-IP in BTSC168 cells transduced with EV or FLAG-ZBTB18. (**D**) Quantification of co-IP results. (**E-F**) qPCR showing binding of LSD1 (**E**) and ZNF217 (**F**) at selected SREBP gene promoters in SNB19 cells transduced with EV, ZBTB18 and ZBTB18-mut. ZNF217 and LSD1 binding increase in presence of ZBTB18. Graphs show representative qPCR results (n=3 technical replicates) of three biological replicates and are expressed in % input as indicated. (**G-I**) qPCR showing binding of LSD1 (**G**), H3K4me2 (**H**) and H3K9me2 (**I**) at selected SREBP gene promoters in BTSC168 cells transduced with EV, ZBTB18 and ZBTB18-mut. Graphs show representative qPCR results (n=3 technical replicates) of at least two biological replicates and are expressed in % input as indicated.

### ZBTB18 inhibits LSD1 demethylation activity to promote its scaffolding function

While ZBTB18-mediated increase of the H3K9me2 repressive marker at the SREBP gene promoter could be an indirect effect due to the recruitment/activation of histone methylase activities, the inhibition of H3K4 methylation activity by LSD1 appeared paradoxical considering LSD1 known repressor role. Therefore, we analyzed the function of LSD1 in more detail by performing CRISPR/Cas9 knockdown in the GBM stem cells-like culture GBM#22 by using two specific gRNAs (Figure 6A). Interestingly, the majority of the genes tested were downregulated upon LSD1 knockdown (Figure 6B) suggesting that LSD1 positively regulates SREBP gene expression. We performed similar experiments using a short hairpin RNA directed to LSD1 in GBM#22 and BTSC475 (Figure 6C-F); here, SREBF1 and SCD were confirmed to be the major LSD1 targets. ChIP analysis of H3K4me2 and H3K9me2 marks at the promoter of SREBP genes upon LSD1 knockdown showed enrichment for both histone marks (Figure 6G). This indicates that LSD1 activation of SREBP genes is accompanied by demethylation of both H3K4me2 and H3K9me2, which could also be an indirect effect due to the recruitment of histone demethylases. Similarly, treatment with the LSD1 inhibitor RN1 in BTSC168 caused enrichment of H3K4me2 and H3K9me2 marks at selected SREBP gene promoters (Figure 6H-I). Therefore, the observation that ZBTB18 expression augments H3K4me2 and H3K9me2 levels is consistent with the hypothesis that ZBTB18 inhibits LSD1 activating function. Interestingly, RN1 treatment in BTSC168 and BTSC268 favored LSD1-ZNF217 and CTBP interaction (Figure 6J-K) prompting us to hypothesize that, inhibition of LSD1 demethylase activity favors its scaffolding function and the interaction with other corepressors (i.e. ZNF217). With a similar mechanism, ZBTB18 expression facilitates the interaction of the ZNF217 corepressor with CTBP2, as well as ZNF217 enrichment at the SREBP gene promoters (Figure 4). Therefore, LSD1 scaffolding function could be implicated in the mechanism of ZBTB18-mediated repression. In conclusion, we propose that ZBTB18 represses SREBP gene expression by 1) inhibiting CTBP and LSD1-mediated transcriptional activation and 2) promoting that assembly of a LSD1-CTBP-ZNF217 repressive complex that seem not to require LSD1-demethylse activity. In the future, more studies to investigate the role of all the CTBP-LSD1 complex components (i.e. ZNF217 and HDAC) in fatty acid synthesis regulation will be required to better define new potential targets in GBM.

**Figure 6.**
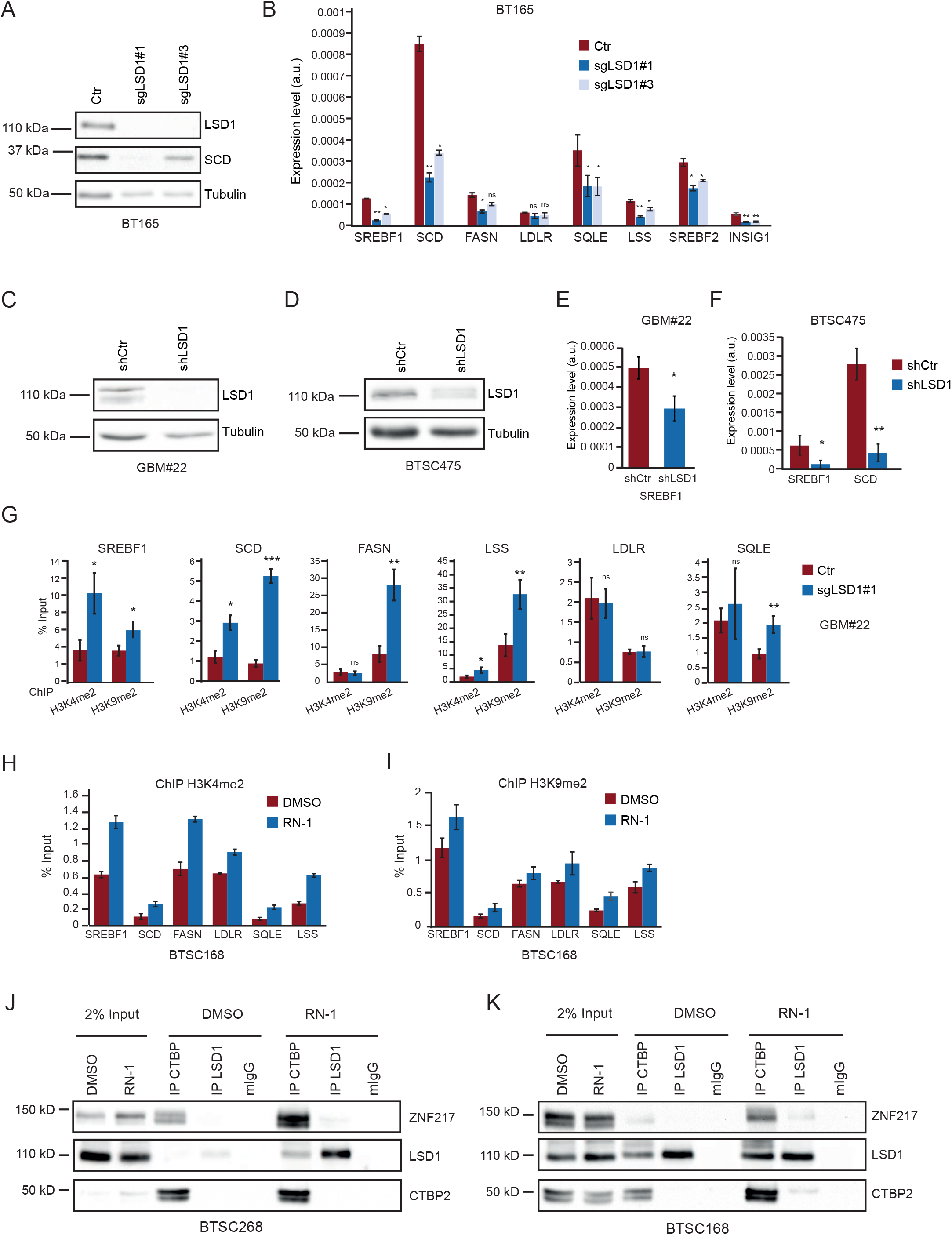
LSD1 functions as activator of SREBP genes independent from its H3K4 and H3K9 demethylase activity. (**A**) WB analysis of LSD1 and SCD expression in GBM#22 cells upon CRISPR/Cas9-mediated LSD1 knockdown with two sgRNA. (**B**) qPCR showing expression levels of selected SREBP genes upon LSD1 depletion. n=3 biological replicates; error bars ± s.d. *p < 0.05, **p < 0.01, ***p < 0.001 by Student’s t-test. Gene expression was normalized to 18s RNA. (**C-D**) WB analysis of LSD1 expression in GBM#22 (**C**) and BTSC475 cells (**D**) transduced with shLSD1. (**E-F**) qPCR analysis of SREBF1 and SCD expression in GBM#22 (**E**) and BTSC475 (**F**) cells upon shRNA-mediated LSD1 silencing. n=3 biological replicates; error bars ± s.d. *p < 0.05, **p < 0.01, ***p < 0.001 by Student’s t-test. Gene expression was normalized to 18s RNA. (**G**) qPCR showing enrichment of H3K4me2 and H3K9me2 marks at the promoter of selected SREBP genes, upon LSD1 knockout in GBM#22 cells. n=3 biological replicates; error bars ± s.d. *p < 0.05, **p < 0.01, ***p < 0.001 by Student’s t-test. Results are expressed in % input as indicated. (**H-I**) qPCR showing enrichment of H3K4me2 (**H**) and H3K9me2 (**I**) marks at the promoter of selected SREBP genes, upon treatment with RN-1. Graphs show representative qPCR results (n=3 technical replicates) of three biological replicates and are expressed in % input as indicated. (**J-K**) WB analysis of LSD1 or CTBP co-IP in BTSC268 (**J**) and BTSC168 (**K**) cells upon treatment with RN-1.

## Discussion

Here, we have identified a new role of ZBTB18, CTBP and LSD1 in the regulation of fatty acids synthesis, which is considered a hallmark of cancer, including GBM. We show that the tumor suppressor ZBTB18 interacts with the cofactors CTBP1/2 and represses the expression of SREBP genes, involved in *de novo* lipogenesis. CTBP is known to be associate in protein complexes containing the histone demethylase LSD1. We show that in GBM cells, which express no or low levels of ZBTB18, CTBP and LSD1 activate the expression of SREBP genes; however, when ectopic expressed, ZBTB18 leads to SREBP gene repression by inhibiting CTBP-associated complex activity. Consistent with such a new function, ZBTB18 expression is paired with the reduction of several phospholipid species, which rely on fatty acids availability to be assembled, and with decreased lipid droplet content inside the cells (Figure 7).

**Figure 7.**
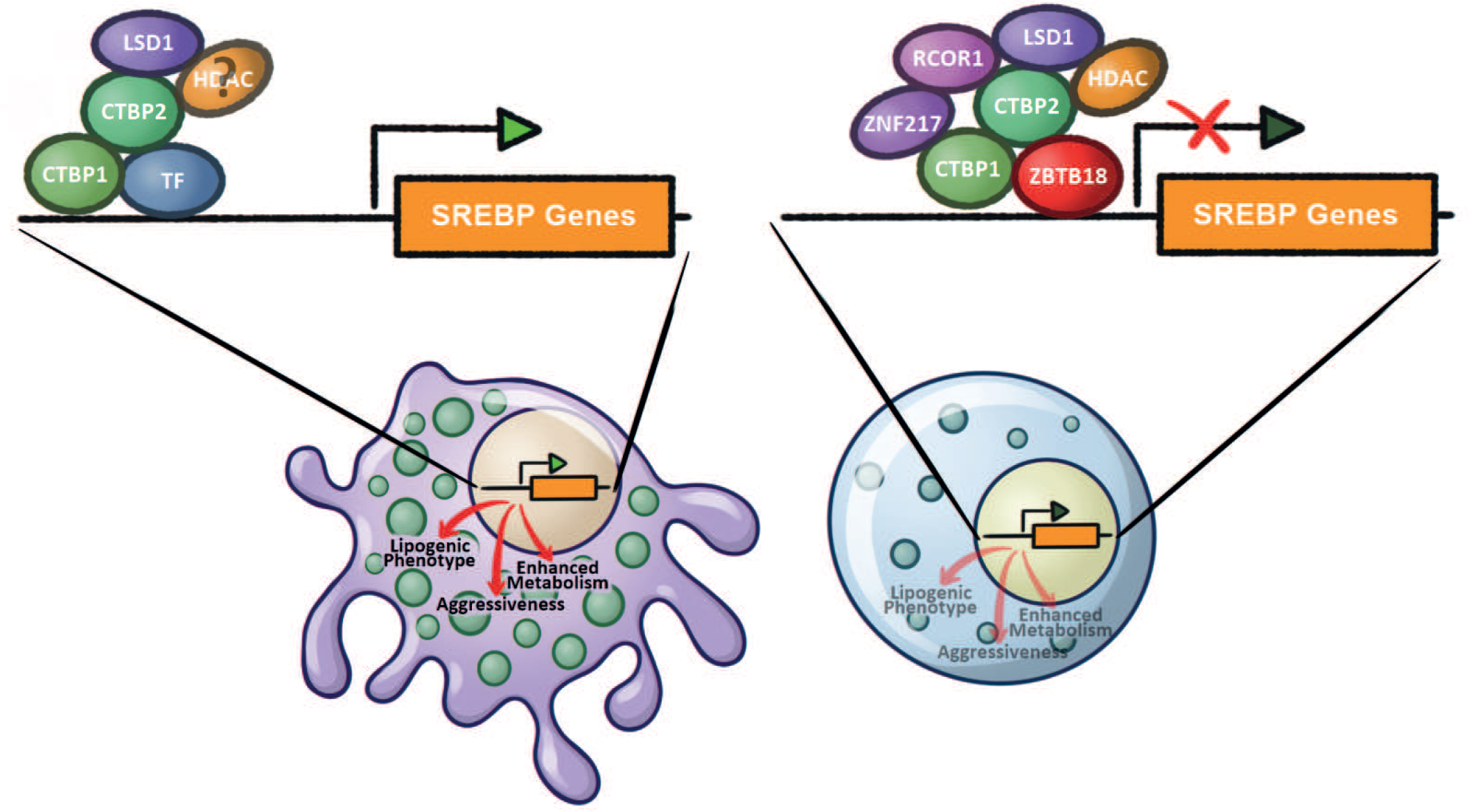
Model of ZBTB18 and CTBP2-mediated regulation of SREBP genes expression. In absence of ZBTB18, CTBP and LSD1 activate the expression of SREBP genes. Other TFs or cofactors could be involved. When expressed, ZBTB18 inhibits LSD1 demethylase activity and recruits a repressive complex at the promoter of SREBP genes.

### CTBPs act as transcriptional activators of SREBP genes

The interaction between ZBTB18 and CTBP2 initially prompted us to consider CTBP2 as a putative ZBTB18 corepressor. However, genome-wide gene expression analysis indicated that ZBTB18 and CTBP2 play opposite roles in the transcriptional regulation of a common set of genes. Among them, EMT and SREBP genes are highly represented and appeared to be activated by CTBP2 and repressed by ZBTB18. Many studies focusing on cell differentiation have shown that CTBP2 can prime its position in actively transcribed genes and that its binding to different transcription factors can cause the switch to a repressive state (Boxer et al. 2014; Kim et al. 2015). In addition, it has been shown that CTBP can be part of activator complexes; for example, CTBP2 and its associated proteins, LSD1 and NCOR1, have been implicated in transcriptional activation by the basic helix-loop-helix transcription factor NeuroD1, in gastrointestinal endocrine cells (Ray et al. 2014). Moreover, CTBP2 has been reported to act as a co-activator of retinoic acid receptor (RAR) (Bajpe et al. 2013).

### ZBTB18 could regulate mesenchymal transformation through SREBP gene regulation

We previously demonstrated that ZBTB18 functions as a repressor of mesenchymal genes, which is expressed at low levels in glioblastoma, mostly due to promoter methylation (Fedele et al. 2017). Therefore, the characterization of ZBTB18 activity in glioblastoma has required overexpression studies. Here we have attempted to knockdown ZBTB18 in BTSC475, a primary GBM cell line that shows some basal level of ZBTB18. Although our results suggest that ZBTB18 loss leads to lipid droplets accumulation, the level of SREBP gene re-expression is modest, suggesting that other mechanisms could be involved. Moreover, CTBP and LSD1 knockout in BTSC475 cells results in the downregulation of SREBP genes, suggesting that here CTBP and LSD1 are mostly participating in gene activation and that due to its low expression ZBTB18 is not sufficient to build a repressive complex on SREBP gene promoters. Interestingly, SREBPs have been recently implicated in mesenchymal transition, in glioblastoma, breast cancer and endothelial cells (Zhang et al. 2019; Martin et al. 2021; Schmitt et al. 2021). Therefore, repressing SREBP-regulated genes could be an additional mechanism through which ZBTB18 counteracts mesenchymal transformation in non-mesenchymal gliomas. Since ZBTB18 is more expressed in low grade gliomas, knockout assays should be established in such cell lines. However, the limitation of low grade gliomas-derived cells has so far represented a major obstacle for the proper establishment of these studies.

### LSD1 plays a role as activator of SREBP genes

Our further analysis suggests that ZBTB18 could interfere with CTBP transcriptional activation of SREBP genes by inhibiting the enzymatic activity of its associated complex, including LSD1-mediated demethylation of H3K4me2 and H3K9me2. While the majority of the studies indicate that LSD1 demethylates mono and di-methylated H3K4, its role as H3K9me2 demethylase has also been reported, mostly in association with nuclear receptors (Metzger et al. 2005; Garcia-Bassets et al. 2007). Although it cannot be excluded that increased levels of H3K9me2 upon ZBTB18 expression could result from other corepressor activities (i.e. specific histone methyltransferases), our data using both LSD1 knockdown and the inhibitor RN1 indicate that, in GBM cells, LSD1 also displays a H3K9me2 demethylase activity, which is important for LSD1-mediated expression of SCD and SREBF1. Interestingly, both positive and negative changes of H3K9me2 have been implicated in the development and progression of various types of cancer due to its influence on cell differentiation, apoptosis, and treatment response (Schulte et al. 2009; Chen et al. 2015). Thus, the role of H3K9me2 in promoting or restringing cancer might depend on the genomic context as well as on the activity of specific histone methylation and demethylation enzymes. The observation that LSD1 knockout and inactivation cause increase of both dimethyl H3K4 and H3K9 is in line with previous findings by Garcia-Bassets and colleagues who demonstrated that both H3K4me2 and H3K9me2 marks are simultaneously decreased upon ER and LSD1 dependent gene activation (Garcia-Bassets et al. 2007). Interestingly, the author also showed that LSD1 contributes to other TFs-mediated activation namely NFκB, AP-1 and βRAR. Furthermore, our data indicate that, in presence of ZBTB18, LSD1 and CTBP more efficiently interact to each other and to the transcriptional corepressor ZNF217. Similar results were obtained when LSD1 demethylase activity was inhibited by RN1. Therefore, we argue that blocking LSD1 demethylase activity could favor LSD1 stability and scaffolding function. This hypothesis is consistent with recent studies indicating that LSD1 can repress gene expression independently from its histone demethylase activity (Sehrawat et al. 2018; Ravasio et al. 2020). In particular, Sehrawat and colleagues demonstrated that LSD1 gene regulation in prostate cancer is mediated by interaction with ZNF217 independently of its demethylase activity.

### ZBTB18 expression affects the content if phospholipids in the cell

Our lipidomics study shows that expression of ZBTB18 in GBM cells affects the synthesis of phospholipids, in particular those containing unsaturated fatty acids. An elevated phospholipid production rate is a trait of rapidly dividing cells, such as tumor cells, as they are required to form the cellular membranes of the new cells. The reduced amount of several phospholipid species upon ZBTB18 ectopic expression might be linked to the previously observed decreased proliferation rate of these cells (Fedele et al. 2017).

It has been previously reported that malignant glioma tissues contain high levels of polyunsaturated fatty acids (Srivastava et al. 2010). Since the SREBF1 target SCD converts newly synthesized saturated fatty acids to unsaturated fatty acids (Cohen et al. 2002), it is possible that ZBTB18-mediated SCD repression accounts for the observed reduction of unsaturated and polyunsaturated fatty acids.

The abundance of lipid droplets within the cell depends on several factors including *de novo* lipid synthesis, uptake of lipids from the environment, and mobilization rate, according to the lipid demand for cellular functions (Olzmann and Carvalho 2019). Our data suggest that ZBTB18 specifically affects the lipid biosynthesis but not the ability of GBM cells to gather lipids from external sources. In fact, while most of the cells rely on external sources of fatty acids, cancer cells have been shown to be able to synthesize their own lipids and recent studies have provided evidence of a relevant role of fatty acid oxidation on GBM tumor growth (Duman et al. 2019).

In conclusion, we unravel a new epigenetic mechanism of transcriptional regulation employed by the tumor suppressor ZBTB18 to repress SREBP genes and *de novo* lipogenesis in GBM. Epigenetic changes have emerged to be a critical step for tumorigenesis and metastasis but the key epigenomic events in cancer cell transformation still remain poorly understood. Our findings contribute to overcome this gap of knowledge. Due to the role of ZBTB18 as a negative regulator of the mesenchymal transformation in GBM, our results have important implications in cancer therapy as they might help to find more effective strategies to diagnose and treat glioblastoma. Furthermore, given the recognized oncogenic role of CTBP2, LSD1 and ZNF217 in other cancers, their implication in *de novo* lipogenesis could have a broader impact in cancer research.

## Materials and methods

### Cell culture

SNB19, LN229 and HEK293T cells were cultured as previously described (Fedele et al. 2017). For lipid starvation, SNB19 cells were grown in DMEM supplemented with 10% lipids depleted FBS (Biowest). Primary glioblastoma stem cells-like BTSC233, BTSC268, BTSC168 and BTSC475 were generated in our laboratory in accordance with an Institutional Review Board-approved protocol (Fedele et al. 2017). BTSC3082 were generated at the University of Uppsala (Xie et al. 2015) and kindly provided by Dr. Nelander. Primary glioblastoma stem cells-like GBM#22 were established at the European Institute of Oncology, Milan, Italy, kindly gifted by Dr. Pelicci. All primary glioblastoma stem cells-like were grown Neurobasal medium (Life Technologies) containing B27 supplement (Life Technologies), FGF (20 ng/ml, R&D Systems), EGF (20 ng/ml, R&D Systems), (LIF 20 ng/ml, Genaxxon Biosciences), Heparin (2 μg/ml, Sigma), and glutamax (Invitrogen). BTSC3082 were grown adherent on dishes previously coated with laminin (Life Technologies). All cells were mycoplasma-free. SNB19 LN229 and HEK293T cells have been authenticated on 3/2/2017 (SNB19 and LN229) and on 11/26/19 (SNB19 and HEK293T) by PCR-single-locus-technology (Eurofins Medigenomix). For CTBP1/2 inhibition, SNB19 and BTSC 233 cells were treated with 10 mM 4-methylthio-2-oxobutyric acid (MTOB, Sigma) for 24h. To inhibit LSD1, GBM cells were treated with 5µM of RN1 (Sigma Aldrich) for 96h.

### RNA extraction and quantitative real time-PCR

Total RNA was extracted from cell culture using miRNeasy Mini Kit (Qiagen) according to the manufacturer’s instructions. First strand cDNA synthesis was generated using the Superscript III First-Strand Synthesis System for RT-PCR (Life Technologies) following the manufacture’s protocol. Quantitative RT-PCR was performed using Kapa SYBR Fast (Sigma). SREBP genes primer sequences are listed in Supplementary Table S3. Primers for SERPINE1, TNFAIP6, CD97, LGALS1, S100A6 and ID1 were previously described (Fedele et al. 2017).

### Gene expression analysis

For gene expression analysis, 1.5 μg of total RNA was processed and analyzed at the DKFZ in Heidelberg (Germany). Hybridization was carried out on Illumina HumanHT-12v4 expression BeadChip. Microarray data were further analyzed by gene set enrichment analysis (GSEA) (http://www.broadinstitute.org/gsea/index.jsp). Microarray gene accession number: GSE138890. Differentially regulated genes were identified using the limma R package (Ritchie et al. 2015). Genes with adjusted pvalue < 0.05 and absolute fold-change > 0.5 were considered as significantly regulated. A gene-sets enrichment analysis was performed using Fisher’s exact test on the Consensus Path DB (Kamburov et al. 2013), comparing these selected genes against the whole set of quantified genes. Significance threshold was set to adjusted pvalue < 0.05. Gene correlation, SREBF1 gene expression and patient survival analysis was performed using the GlioVis platform (Bowman et al. 2017) and data from The Cancer Genome Atlas (TCGA) (https://www.cancer.gov/tcga) and Chinese Glioma Genome Atlas (CGGA) project (Zhao et al. 2017).

### Immunoblotting

Total protein extracts were prepared as previously described (Fedele et al. 2017). Western blots were performed using the antibodies listed in Supplementary Table S5.

### Co-immunoprecipitation and mass spectrometry and SILAC

For co-immunoprecipitation (co-IP) GBM cells were lysed in 1 ml of co-IP buffer (150 mM NaCl, 50 mM TrisHCl pH 7.5, 10% glycerol, 0.2% Igepal, 1mM EDTA) supplemented with Halt^TM^ protease and phosphatase inhibitor cocktail (1mM, Thermo Scientific), and PMSF (1mM, Sigma),vortexed for 30 seconds and kept on ice for 30 minutes. After centrifugation at 13,200 rpm for 20 minutes at 4°C a pre-clearing step was performed by adding 25 µl of Protein A/G PLUS-Agarose beads (Santa Cruz) and incubating the samples for 1 hour at 4°C in rotation. The recovered supernatant was incubated overnight with the antibodies listed in Supplementary Table S6. 20 µl of lysate (input) were removed and mixed with equal volume of 2X laemmli buffer for western blot analysis. 20 µl of equilibrated protein A/G beads were added to each lysates and incubated for 2 hours at 4°C. The beads were then washed for 4 times with 1 ml of co-IP buffer before eluting the co-precipitated proteins with 20 µl of 2X laemmli buffer. Samples were stored at −20°C or directly analyzed by western blot. For ZBTB18 co-IP in BTSC3082 and BTSC268 cells,lysates were incubated overnight with rabbit anti-ZBTB18 (Proteintech #12714-1-AP). For mass spectrometry (MS) analysis, the eluted co-precipitated proteins were separated by gel electrophoresis and gel fragments were processed as described in (Tucci et al. 2018) and then measured on a LTQ Orbitrap XL (Thermo Scientific) instrument. Data acquisition and analysis was performed using Xcalibur (Thermo Scientific, Germany) and MaxQuant 1.4.1.2 softwares. To run SILAC, SNB19 cells were SILAC labelled using Arg0/Lys0 (low, L) and Arg10/Lys8 (high, H) (Silantes). Afterwards cells were transduced with pCHMWS-EV (L) or pCHMWS-FLAG-ZBTB18 (H) lentiviral particles. Transduced SNB19 were mixed and lysed as described above. Total protein extracts were incubated with rabbit anti-CTBP2 (Cell Signaling #13256S) and the precipitated fraction subjected to MS to identify interacting proteins and calculate H/L ratios. All measurements and analysis were performed at the Core Facility Proteomics of the Center for Biological System Analysis (ZBSA) at the University of Freiburg, Germany. MS analysis of ZBTB18 co-IP in BTSC268 was perfomed at the Proteomic Platform – Core Facility of the Freiburg Medical Center. A detailed protocol of the procedure is included in the supplementary materials section.

The protein interactions from this publication have been submitted to the IMEx (www.imexconsortium.org) consortium through IntAct (Orchard et al. 2014) and assigned the identification number IM-27496.

### Migration, proliferation and apoptosis assay

Migration, proliferation and apoptosis assay were performed as previously described (Fedele et al. 2017). For apoptosis assay, transduced SNB19 cells were seeded in a 96 well (dark plate) at a density of 1×10^3^ per well. The following day activity of caspases 3 and 7 was measured using the Caspase-Glo® 3/7 Assay (Promega), according to the manufacturer’s instruction.

### LSD1 activation assay

For LSD1 activity assay, SNB19 cells were transduced with EV, ZBTB18 or ZBTB18-mut as described above or treated with 5 µM RN-1 inhibitor (Calbiochem), for 96h. Nuclear lysates were prepared with EpiQuik™ Nuclear Extraction Kit (Epigenetk). Measurement of LSD1 activity was performed with Epigenase LSD1 Demethylase Activity/Inhibition Assay Kit (Epigentek) according to the manufacturer’s instructions. LSD1 activity was quantified by Infinite 200 PRO Spectrophotometer (Tecan) at 450 nm and 655 nm, following the manufacturer’s instructions.

### ChIP-seq and quantitative ChIP

For ChIP-seq, SNB19 cells were transduced and incubated with ChIP Crosslink Gold solution (Diagenode) and processed using the iDeal ChIP-seq kit for Transcription Factors (Diagenode), according to the manufacturer’s instruction. Antibodies are listed in table S7. Libraries preparation and ChIP-seq were performed at Diagenode (https://www.diagenode.com). ChIPseq gene accession number: GSE140002. Downstream analyses of were executed with R (3.6.0). For each condition two independent ChIP were performed. Venn analysis was performed with the ChIPpeakAnno R package (Zhu et al. 2010). A single base overlap threshold was used to identify the common peaks between the 3 conditions. Peaks were divided according to the Venn subgroups and annotated with the nearest gene using ChIPseeker (Yu et al. 2015) and TxDb.Hsapiens UCSC.hg19.knownGene R packages. Enrichment analysis of gene-regions was done via a Fisher’s exact test using 100000 regions of 200bp, randomly selected on the human genome, as background. ReMap (Cheneby et al. 2019) was used to look at overlap with other TFs and cofactors peaks from published data in other cell lines. The statistical significance of the overlap was calculated by repeating the analysis using the same number of randomly selected regions of 2000 bp 1000 time. The p pvalue was calculated from the empirical cumulative distribution. Motives enrichment analysis was performed with the Homer software (Heinz et al. 2010). A detailed description of quantitative ChIP is included in the supplementary materials section. Primers are listed in SupplementaryTable S4.

### Lipidomics Analysis

For lipid profiling, cells were washed, quenched and lysed as previously described (Lagies et al. 2018) and lipids were extracted as described by (Sapcariu et al. 2014). Internal standard containing organic phase was evaporated to dryness and reconstituted in isopropanol:acetonitrile:water 2:1:1 and subjected to targeted LC-MS lipid profiling. A BEH C18 2.1 mm x 100 mm, 1.8 µm column (Waters Corporation) was used with the following chromatographic program: 40 % solvent B (10 mM NH_4_CHO_2_, 0.1% formic acid, isopropanol:acetonitrile 9:1) was hold for 2 minutes, to 98% B within 10 minutes, hold for 2 minutes, to 40% B within 0.5 minutes and hold for 5.5 minutes. Solvent A was 10 mM NH_4_CHO_2_, 0.1% formic acid, acetonitrile:water 3:2. The flow rate was set to 300 µl/minute and the column temperature was 55°C. The method was further verified by analyzing lipid standards with the same chromatography coupled to a high resolution mass spectrometer (Synapt G2Si, Waters Corporation). Samples were kept at 12°C and injected in randomized order with pooled quality control samples injected at regular intervals. Lipids were monitored by MRM and SIM scan modes using an Agilent 6460 triple quadrupole. Raw data were analyzed by Agilent Quantitative Analysis software. Samples were normalized to internal standard, QC-filtered and range-scaled for principal component analysis and heat map generation. Significance was determined by ANOVA including FDR multiple testing correction (q-value cut off: 0.05). Statistical analyses and visualization were carried out by MetaboAnalyst 4.0 (Chong et al. 2018).

### Immunostaining and lipid staining

SNB19 cells were grown in 4-well chamber slides either with 10% FBS DMEM or 10% lipid-depleted FBS (Biowest) DMEM for 48 hours. For lipid starvation cells were kept for additional 2 hours either in 10% lipid-depleted FBS DMEM or in 10% FBS DMEM. All cells were then fixed with 4% paraformaldehyde in PBS and processed for the staining. ZBTB18 was labelled with anti-ZBTB18 rabbit polyclonal primary antibody (Proteintech #12714-1-AP) and anti-Rabbit IgG (H+L) Alexa Fluor 647 (goat, Thermo Fisher Scientific) secondary antibody. Lipid droplets were stained with 0.5 µg/ml Bodipy TMR-X SE (Thermo Fisher Scientific) in 150 mM NaCl for 10 minutes at room temperature. Nuclei were counterstained with 4′,6-diamidino-2-phenylindole (DAPI, Sigma-Aldrich). Pictures were acquired using a FSL confocal microscope (Olympus).

### Lipid uptake

SNB19 cells were seeded in 4-well chamber slides and infected as described above. The cells were then incubated with 50 nM Bodipy-C16 (Thermo Fisher Scientific) in PBS for 15 minutes at room temperature. Subsequently, the samples were fixed with 4% paraformaldehyde in PBS, counterstained with DAPI and imaged using a FSL confocal microscope (Olympus).

### Statistical analysis

For the statistical analysis of the RT-qPCR data are judged to be statistically significant when p < 0.05 by two-tailed Student’s t test. The number of replicates and the definition of biological versus technical replicates is indicated in each Figure legends.

## Acknowledgments

We thank Mahmoud Abdelkarim, Jonathan Goldner, Verena Haase, Marzieh Mohammadi and Jutith Treiber for technical assistance. We thank Veronica Dumit and Lisa Winner (University of Freiburg, ZBSA) for MS analysis and Marc Timmers for advice and discussion. We are grateful to M. Squatrito (CNIO, Madrid, Spain) for sharing protocols and advice on CRISPR/Cas9 methodology. The results shown here are in part based upon data generated by the TCGA Research Network: https://www.cancer.gov/tcga. This work was financed by the German’s Cancer Aid grant (Deutsche Krebshilfe, 70113120) to MSC. AI was supported by the Research Committee (Forschungskomission) of the Faculty of Medicine, University of Freiburg. The Proteomic Platform–Core Facility was supported by the Research Committee (Forschungskomission) of the Faculty of Medicine, University of Freiburg.

## Author contribution

RF, AI, JPB, APM, AW, SY, RS, VR and EK performed experiments. GA and MB performed microarray and ChIP-seq analysis. SL and BK performed lipidomics analysis. DO, SF and GP provided LSD1 KO cells and LSD1 silencing reagents. SGT performed MS analysis. DHH perfomed in silico analysis. MP provided reagents. MSC conceived and supervised the study. MSC, RF and AI wrote the manuscripts.

## References

Achouri Y, Noel G, Van Schaftingen E. 2007. 2-Keto-4-methylthiobutyrate, an intermediate in the methionine salvage pathway, is a good substrate for CtBP1. Biochemical and biophysical research communications 352: 903–906.

Bajpe PK, Heynen GJ, Mittempergher L, Grernrum W, de Rink IA, Nijkamp W, Beijersbergen RL, Bernards R, Huang S. 2013. The corepressor CTBP2 is a coactivator of retinoic acid receptor/retinoid X receptor in retinoic acid signaling. Molecular and cellular biology 33: 3343–3353.

Bensaad K, Favaro E, Lewis CA, Peck B, Lord S, Collins JM, Pinnick KE, Wigfield S, Buffa FM, Li JL et al. 2014. Fatty acid uptake and lipid storage induced by HIF-1alpha contribute to cell growth and survival after hypoxia-reoxygenation. Cell reports 9: 349–365.

Bhat KPL, Balasubramaniyan V, Vaillant B, Ezhilarasan R, Hummelink K, Hollingsworth F, Wani K, Heathcock L, James JD, Goodman LD et al. 2013. Mesenchymal differentiation mediated by NF-kappaB promotes radiation resistance in glioblastoma. Cancer Cell 24: 331–346.

Bowman RL, Wang Q, Carro A, Verhaak RG, Squatrito M. 2017. GlioVis data portal for visualization and analysis of brain tumor expression datasets. Neuro-oncology 19: 139–141.

Boxer LD, Barajas B, Tao S, Zhang J, Khavari PA. 2014. ZNF750 interacts with KLF4 and RCOR1, KDM1A, and CTBP1/2 chromatin regulators to repress epidermal progenitor genes and induce differentiation genes. Genes & development 28: 2013–2026.

Carro MS, Lim WK, Alvarez MJ, Bollo RJ, Zhao X, Snyder EY, Sulman EP, Anne SL, Doetsch F, Colman H et al. 2010. The transcriptional network for mesenchymal transformation of brain tumours. Nature 463: 318–325.

Chen M, Zhu N, Liu X, Laurent B, Tang Z, Eng R, Shi Y, Armstrong SA, Roeder RG. 2015. JMJD1C is required for the survival of acute myeloid leukemia by functioning as a coactivator for key transcription factors. Genes & development 29: 2123–2139.

Cheneby J, Menetrier Z, Mestdagh M, Rosnet T, Douida A, Rhalloussi W, Bergon A, Lopez F, Ballester B. 2019. ReMap 2020: a database of regulatory regions from an integrative analysis of Human and Arabidopsis DNA-binding sequencing experiments. Nucleic Acids Res.

Cheng C, Ru P, Geng F, Liu J, Yoo JY, Wu X, Cheng X, Euthine V, Hu P, Guo JY et al. 2015. Glucose-Mediated N-glycosylation of SCAP Is Essential for SREBP-1 Activation and Tumor Growth. Cancer Cell 28: 569–581.

Chong J, Soufan O, Li C, Caraus I, Li S, Bourque G, Wishart DS, Xia J. 2018. MetaboAnalyst 4.0: towards more transparent and integrative metabolomics analysis. Nucleic Acids Res 46: W486–W494.

Cohen P, Miyazaki M, Socci ND, Hagge-Greenberg A, Liedtke W, Soukas AA, Sharma R, Hudgins LC, Ntambi JM, Friedman JM. 2002. Role for stearoyl-CoA desaturase-1 in leptin-mediated weight loss. Science 297: 240–243.

Di LJ, Byun JS, Wong MM, Wakano C, Taylor T, Bilke S, Baek S, Hunter K, Yang H, Lee M et al. 2013. Genome-wide profiles of CtBP link metabolism with genome stability and epithelial reprogramming in breast cancer. Nature communications 4: 1449.

Duman C, Yaqubi K, Hoffmann A, Acikgoz AA, Korshunov A, Bendszus M, Herold-Mende C, Liu HK, Alfonso J. 2019. Acyl-CoA-Binding Protein Drives Glioblastoma Tumorigenesis by Sustaining Fatty Acid Oxidation. Cell Metab 30: 274–289 e275.

Fedele M, Cerchia L, Pegoraro S, Sgarra R, Manfioletti G. 2019. Proneural-Mesenchymal Transition: Phenotypic Plasticity to Acquire Multitherapy Resistance in Glioblastoma. International journal of molecular sciences 20.

Fedele V, Dai F, Masilamani A, Heiland DH, Kling E, Gatjens-Sanchez A, Ferrarese R, Platania L, Doostkam S, Kim H et al. 2017. Epigenetic Regulation of ZBTB18 Promotes Glioblastoma Progression. Mol Cancer Res.

Garcia-Bassets I, Kwon YS, Telese F, Prefontaine GG, Hutt KR, Cheng CS, Ju BG, Ohgi KA, Wang J, Escoubet-Lozach L et al. 2007. Histone methylation-dependent mechanisms impose ligand dependency for gene activation by nuclear receptors. Cell 128: 505–518.

Geng F, Cheng X, Wu X, Yoo JY, Cheng C, Guo JY, Mo X, Ru P, Hurwitz B, Kim SH et al. 2016. Inhibition of SOAT1 Suppresses Glioblastoma Growth via Blocking SREBP-1-Mediated Lipogenesis. Clinical cancer research : an official journal of the American Association for Cancer Research 22: 5337–5348.

Goldmann T, Zeller N, Raasch J, Kierdorf K, Frenzel K, Ketscher L, Basters A, Staszewski O, Brendecke SM, Spiess A et al. 2015. USP18 lack in microglia causes destructive interferonopathy of the mouse brain. The EMBO journal 34: 1612–1629.

Guo D, Reinitz F, Youssef M, Hong C, Nathanson D, Akhavan D, Kuga D, Amzajerdi AN, Soto H, Zhu S et al. 2011. An LXR agonist promotes glioblastoma cell death through inhibition of an EGFR/AKT/SREBP-1/LDLR-dependent pathway. Cancer discovery 1: 442–456.

Heinz S, Benner C, Spann N, Bertolino E, Lin YC, Laslo P, Cheng JX, Murre C, Singh H, Glass CK. 2010. Simple combinations of lineage-determining transcription factors prime cis-regulatory elements required for macrophage and B cell identities. Molecular cell 38: 576–589.

Horton JD, Goldstein JL, Brown MS. 2002. SREBPs: activators of the complete program of cholesterol and fatty acid synthesis in the liver. The Journal of clinical investigation 109: 1125–1131.

Kamburov A, Stelzl U, Lehrach H, Herwig R. 2013. The ConsensusPathDB interaction database: 2013 update. Nucleic Acids Res 41: D793–800.

Kim TW, Kang BH, Jang H, Kwak S, Shin J, Kim H, Lee SE, Lee SM, Lee JH, Kim JH et al. 2015. Ctbp2 Modulates NuRD-Mediated Deacetylation of H3K27 and Facilitates PRC2-Mediated H3K27me3 in Active Embryonic Stem Cell Genes During Exit from Pluripotency. Stem Cells 33: 2442–2455.

Lagies S, Pichler R, Kaminski MM, Schlimpert M, Walz G, Lienkamp SS, Kammerer B. 2018. Metabolic characterization of directly reprogrammed renal tubular epithelial cells (iRECs). Scientific reports 8: 3878.

Lewis CA, Brault C, Peck B, Bensaad K, Griffiths B, Mitter R, Chakravarty P, East P, Dankworth B, Alibhai D et al. 2015. SREBP maintains lipid biosynthesis and viability of cancer cells under lipid- and oxygen-deprived conditions and defines a gene signature associated with poor survival in glioblastoma multiforme. Oncogene 34: 5128–5140.

Martin M, Zhang J, Miao Y, He M, Kang J, Huang HY, Chou CH, Huang TS, Hong HC, Su SH et al. 2021. Role of endothelial cells in pulmonary fibrosis via SREBP2 activation. JCI insight 6.

Menendez JA, Lupu R. 2007. Fatty acid synthase and the lipogenic phenotype in cancer pathogenesis. Nature reviews Cancer 7: 763–777.

Metzger E, Wissmann M, Yin N, Muller JM, Schneider R, Peters AH, Gunther T, Buettner R, Schule R. 2005. LSD1 demethylates repressive histone marks to promote androgen-receptor-dependent transcription. Nature 437: 436–439.

Nibu Y, Levine MS. 2001. CtBP-dependent activities of the short-range Giant repressor in the Drosophila embryo. Proceedings of the National Academy of Sciences of the United States of America 98: 6204–6208.

Olzmann JA, Carvalho P. 2019. Dynamics and functions of lipid droplets. Nature reviews Molecular cell biology 20: 137–155.

Orchard S, Ammari M, Aranda B, Breuza L, Briganti L, Broackes-Carter F, Campbell NH, Chavali G, Chen C, del-Toro N et al. 2014. The MIntAct project--IntAct as a common curation platform for 11 molecular interaction databases. Nucleic Acids Res 42: D358–363.

Ostrom QT, Gittleman H, Fulop J, Liu M, Blanda R, Kromer C, Wolinsky Y, Kruchko C, Barnholtz-Sloan JS. 2015. CBTRUS Statistical Report: Primary Brain and Central Nervous System Tumors Diagnosed in the United States in 2008-2012. Neuro-oncology 17 Suppl 4: iv1–iv62.

Phillips HS, Kharbanda S, Chen R, Forrest WF, Soriano RH, Wu TD, Misra A, Nigro JM, Colman H, Soroceanu L et al. 2006. Molecular subclasses of high-grade glioma predict prognosis, delineate a pattern of disease progression, and resemble stages in neurogenesis. Cancer Cell 9: 157–173.

Ravasio R, Ceccacci E, Nicosia L, Hosseini A, Rossi PL, Barozzi I, Fornasari L, Zuffo RD, Valente S, Fioravanti R et al. 2020. Targeting the scaffolding role of LSD1 (KDM1A) poises acute myeloid leukemia cells for retinoic acid-induced differentiation. Sci Adv 6: eaax2746.

Ray SK, Li HJ, Metzger E, Schule R, Leiter AB. 2014. CtBP and associated LSD1 are required for transcriptional activation by NeuroD1 in gastrointestinal endocrine cells. Molecular and cellular biology 34: 2308–2317.

Ritchie ME, Phipson B, Wu D, Hu Y, Law CW, Shi W, Smyth GK. 2015. limma powers differential expression analyses for RNA-sequencing and microarray studies. Nucleic Acids Res 43: e47.

Romani M, Pistillo MP, Banelli B. 2018. Epigenetic Targeting of Glioblastoma. Front Oncol 8: 448.

Ru P, Hu P, Geng F, Mo X, Cheng C, Yoo JY, Cheng X, Wu X, Guo JY, Nakano I et al. 2016. Feedback Loop Regulation of SCAP/SREBP-1 by miR-29 Modulates EGFR Signaling-Driven Glioblastoma Growth. Cell reports 16: 1527–1535.

Sapcariu SC, Kanashova T, Weindl D, Ghelfi J, Dittmar G, Hiller K. 2014. Simultaneous extraction of proteins and metabolites from cells in culture. MethodsX 1: 74–80.

Schmitt MJ, Company C, Dramaretska Y, Barozzi I, Gohrig A, Kertalli S, Grossmann M, Naumann H, Sanchez-Bailon MP, Hulsman D et al. 2021. Phenotypic Mapping of Pathologic Cross-Talk between Glioblastoma and Innate Immune Cells by Synthetic Genetic Tracing. Cancer discovery 11: 754–777.

Schulte JH, Lim S, Schramm A, Friedrichs N, Koster J, Versteeg R, Ora I, Pajtler K, Klein-Hitpass L, Kuhfittig-Kulle S et al. 2009. Lysine-specific demethylase 1 is strongly expressed in poorly differentiated neuroblastoma: implications for therapy. Cancer Res 69: 2065–2071.

Sehrawat A, Gao L, Wang Y, Bankhead A, 3rd, McWeeney SK, King CJ, Schwartzman J, Urrutia J, Bisson WH, Coleman DJ et al. 2018. LSD1 activates a lethal prostate cancer gene network independently of its demethylase function. Proceedings of the National Academy of Sciences of the United States of America 115: E4179–E4188.

Shi Y, Lan F, Matson C, Mulligan P, Whetstine JR, Cole PA, Casero RA, Shi Y. 2004. Histone demethylation mediated by the nuclear amine oxidase homolog LSD1. Cell 119: 941–953.

Shi Y, Sawada J, Sui G, Affar el B, Whetstine JR, Lan F, Ogawa H, Luke MP, Nakatani Y, Shi Y. 2003. Coordinated histone modifications mediated by a CtBP co-repressor complex. Nature 422: 735–738.

Srivastava NK, Pradhan S, Gowda GA, Kumar R. 2010. In vitro, high-resolution 1H and 31P NMR based analysis of the lipid components in the tissue, serum, and CSF of the patients with primary brain tumors: one possible diagnostic view. NMR Biomed 23: 113–122.

Tatard VM, Xiang C, Biegel JA, Dahmane N. 2010. ZNF238 Is Expressed in Postmitotic Brain Cells and Inhibits Brain Tumor Growth. Cancer Res.

Tucci S, Mingirulli N, Wehbe Z, Dumit VI, Kirschner J, Spiekerkoetter U. 2018. Mitochondrial fatty acid biosynthesis and muscle fiber plasticity in very long-chain acyl-CoA dehydrogenase-deficient mice. FEBS letters 592: 219–232.

Verhaak RG, Hoadley KA, Purdom E, Wang V, Qi Y, Wilkerson MD, Miller CR, Ding L, Golub T, Mesirov JP et al. 2010. Integrated Genomic Analysis Identifies Clinically Relevant Subtypes of Glioblastoma Characterized by Abnormalities in PDGFRA, IDH1, EGFR, and NF1. Cancer Cell 17: 98–110.

Wang J, Cazzato E, Ladewig E, Frattini V, Rosenbloom DI, Zairis S, Abate F, Liu Z, Elliott O, Shin YJ et al. 2016a. Clonal evolution of glioblastoma under therapy. Nature genetics 48: 768–776.

Wang Q, Hu B, Hu X, Kim H, Squatrito M, Scarpace L, deCarvalho AC, Lyu S, Li P, Li Y, et al. 2017. Tumor Evolution of Glioma-Intrinsic Gene Expression Subtypes Associates with Immunological Changes in the Microenvironment. Cancer Cell 32: 42–56 e46.

Wang Y, Che S, Cai G, He Y, Chen J, Xu W. 2016b. Expression and prognostic significance of CTBP2 in human gliomas. Oncology letters 12: 2429–2434.

Wang Y, Wang H, Zhao Q, Xia Y, Hu X, Guo J. 2015. PD-L1 induces epithelial-to-mesenchymal transition via activating SREBP-1c in renal cell carcinoma. Medical oncology 32: 212.

Xiang C, Frietze KK, Bi Y, Li Y, Dal Pozzo V, Pal S, Alexander N, Baubet V, D’Acunto V, Mason CE et al. 2021. RP58 Represses Transcriptional Programs Linked to Nonneuronal Cell Identity and Glioblastoma Subtypes in Developing Neurons. Molecular and cellular biology 41: e0052620.

Xie Y, Bergstrom T, Jiang Y, Johansson P, Marinescu VD, Lindberg N, Segerman A, Wicher G, Niklasson M, Baskaran S et al. 2015. The Human Glioblastoma Cell Culture Resource: Validated Cell Models Representing All Molecular Subtypes. EBioMedicine 2: 1351–1363.

Yang L, Yang J, Li J, Shen X, Le Y, Zhou C, Wang S, Zhang S, Xu D, Gong Z. 2015. MircoRNA-33a inhibits epithelial-to-mesenchymal transition and metastasis and could be a prognostic marker in non-small cell lung cancer. Scientific reports 5: 13677.

Yu G, Wang LG, He QY. 2015. ChIPseeker: an R/Bioconductor package for ChIP peak annotation, comparison and visualization. Bioinformatics 31: 2382–2383.

Zhang N, Zhang H, Liu Y, Su P, Zhang J, Wang X, Sun M, Chen B, Zhao W, Wang L et al. 2019. SREBP1, targeted by miR-18a-5p, modulates epithelial-mesenchymal transition in breast cancer via forming a co-repressor complex with Snail and HDAC1/2. Cell death and differentiation 26: 843–859.

Zhao Z, Hao D, Wang L, Li J, Meng Y, Li P, Wang Y, Zhang C, Zhou H, Gardner K et al. 2019. CtBP promotes metastasis of breast cancer through repressing cholesterol and activating TGF-beta signaling. Oncogene 38: 2076–2091.

Zhao Z, Meng F, Wang W, Wang Z, Zhang C, Jiang T. 2017. Comprehensive RNA-seq transcriptomic profiling in the malignant progression of gliomas. Sci Data 4: 170024.

Zhu LJ, Gazin C, Lawson ND, Pages H, Lin SM, Lapointe DS, Green MR. 2010. ChIPpeakAnno: a Bioconductor package to annotate ChIP-seq and ChIP-chip data. BMC bioinformatics 11: 237.

