## Supplemental figures and materials for "ZBTB18 inhibits SREBP-dependent fatty acid synthesis by counteracting CTBPs and KDM1A/LSD1 activity in glioblastoma"

### ZBTB18 interacts with CTBP and interferes with CTBP-LSD1 transcriptional activation of SREBP genes in glioblastoma

**Inventory:**

**Figure 1-figure supplement 1**

**Figure 1- figure supplement 2**

**Figure 2- figure supplement 1**

**Figure 2- figure supplement 2**

**Figure 2- figure supplement 3**

**Figure 3- figure supplement 1**

**Figure 4- figure supplement 1**

**Figure 4- figure supplement 2**

**Figure 5- figure supplement 1**

**Supplementary materials.** Includes information on lentiviral vectors, MS and qChIP procedures, shRNAs and gRNA sequences.

**Supplementary table 1**. List of annotated SREBP genes detected by Venn analysis performed on peak coordinates. Peaks at SREBP genes promoter in proximity to the transcription start mostly occur in the “All” group (highlighted in light blue).

**Supplementary tables 2-5.** List of primers used for cloning, site-directed mutagenesis, qRT-PCR and quantitative ChIP.

**Supplementary tables 3-7.** List of antibodies used for western blot, co-IP and ChIP.

Figure 1-figure supplement 1

A

- MS in 3082 (ZBTB18 co-IP with anti ZBTB18 antibody from proteintech, Cte) shows interaction with ZBTB18 and CTBP1

| Gene names | Fasta headers | Number of proteins | Peptides | PEP | Intensity Sample 1 (IgG) | Intensity Sample 2 (ZBTB18 IP-top) | Intensity Sample 3 (ZBTB18-bottom) |
|---|---|---|---|---|---|---|---|
| CTBP1 | >sp\|Q13363-2\|CTBP1_HUMAN Isoform 2 of C-terminal-binding protein 1 OS=Homo sapiens GN=CTBI | 12 | 6 | 1,26E-17 | 0 | 98109000 | 0 |
| ZBTB18 | >sp\|Q99592\|ZBT18_HUMAN Zinc finger and BTB domain-containing protein 18 OS=Homo sapiens GN | 3 | 13 | 2,51E-289 | 0 | 25190000 | 5113200 |

B

- MS in BTSC 268 (ZBTB18 co-IP with anti ZBTB18 antibody from proteintech, Cte) confirms ZBTB18 interaction with CTBP1 and CTBP2

| Protein IDs | Protein names | Gene names | Razor + unique peptides | Unique peptides | Sequence coverage [%] | LFQ intensity BTSC268 IgG | LFQ intensity BTSC268 αZNF238 | Log2 of LFQ intensity BTSC268 IgG | Log2 of LFQ intensity BTSC268 αZNF238 | Log2 Fold Change (FC) of LFQ intensity BTSC268 (αZNF238/IgG control) |
|---|---|---|---|---|---|---|---|---|---|---|
| Q99592;B2RXF5 | Zinc finger and BTB domain-containing protein 18 | ZBTB18 | 7 | 7 | 19,2 | 10000 | 499500000 | 13,2877124 | 28,8959094 | 15,6081971 |
| P56545 | C-terminal-binding protein 2 | CTBP2 | 2 | 2 | 10,1 | 10000 | 10284000 | 13,2877124 | 23,2938982 | 10,0061858 |
| Q13363 | C-terminal-binding protein 1 | CTBP1 | 6 | 4 | 13,2 | 1578300 | 82553000 | 20,58994 | 26,2988173 | 5,70887728 |

C

- SILAC-MS in SNB19 (CTBP2 co-IP with anti CTBP2 antibody) confirms CTBP2 interaction with CTBP1 and ZBTB18

| Protein IDs | Gene names | Number of proteins | Peptides | PEP | Ratio H/L | Ratio H/L normalized | Ratio H/L normalized Sample 1 |
|---|---|---|---|---|---|---|---|
| Q13363;D6RAX2;H0Y8W7;H0Y9M9;H0Y8U5 | CTBP1 | 5 | 15 | 1,39E-254 | 0,39903 | 0,18288 | 0,18288 |
| P56545;Q5SQP8;P56545-2 | CTBP2 | 3 | 17 | 0 | 0,70522 | 0,7909 | 0,7909 |
| Q99592-2;B2RXF5 | ZBTB18 | 2 | 12 | 1,42E-185 | 22,708 | 30,176 | 30,176 |

D

| Gene names | Number of proteins | Peptides |
|---|---|---|
| HSPA5 | 1 | 27 |
| HSPA8 | 16 | 22 |
| CCT8 | 5 | 17 |
| ZBTB18 | 1 | 17 |
| CCT2 | 4 | 16 |
| CCT6A | 7 | 13 |
| CCT5 | 6 | 13 |
| TCP1 | 10 | 12 |
| HSPA9 | 3 | 11 |
| CCT3 | 6 | 9 |
| PCMT1 | 10 | 8 |
| HSPA1A | 4 | 8 |
| EEF1A1P5 | 5 | 8 |
| PRDX1 | 2 | 7 |
| CCT7 | 7 | 7 |
| CCT4 | 4 | 6 |
| PAICS | 4 | 6 |
| HSPA6 | 2 | 6 |
| GSTP1 | 2 | 5 |
| CTBP2 | 3 | 5 |
| FLNA | 4 | 4 |
| CBR1 | 4 | 3 |
| UBB | 29 | 3 |
| CBR3 | 1 | 3 |
| DCD | 2 | 3 |
| CLTC | 3 | 3 |
| CTBP1 | 8 | 3 |
| AMY2B | 5 | 2 |

E

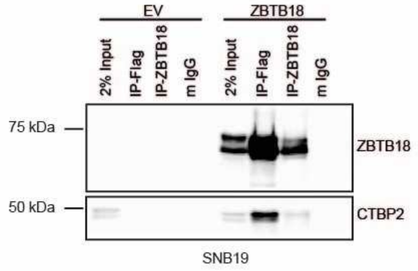


F

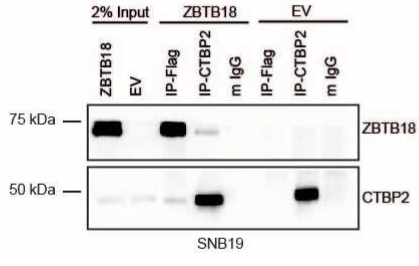

**Figure 1-figure supplement 1. ZBTB18 interacts with CTBP1 and CTBP2 in GBM cells.** (**A-B**) Mass Spec results of FLAG-ZBTB18 co-IP showing detection of CTBP1 (**A**) or CTBP1 and CTBP2 (**B**) in the co-precipitated fraction of BTSC3082 and BTS268 cells respectively. (**C**) SILAC Mass Spec results of CTBP2 co-IP in SNB19 cells transduced with empty vector (EV) or FLAG-ZBTB18, showing detection of FLAG-ZBTB18 in the co-precipitated fraction. (**D**) List of ZBTB18 co-precipitated proteins upon FLAG-ZBTB18 co-IP in SNB19 GBM cells transduced with EV or FLAG-ZBTB18. (**E-F**) WB analysis of FLAG and ZBTB18 co-IPs (**E**) or FLAG and CTBP2 co-IPs (**F**) in SNB19 cells transduced with EV or FLAG-ZBTB18.

Figure 1-figure supplement 2

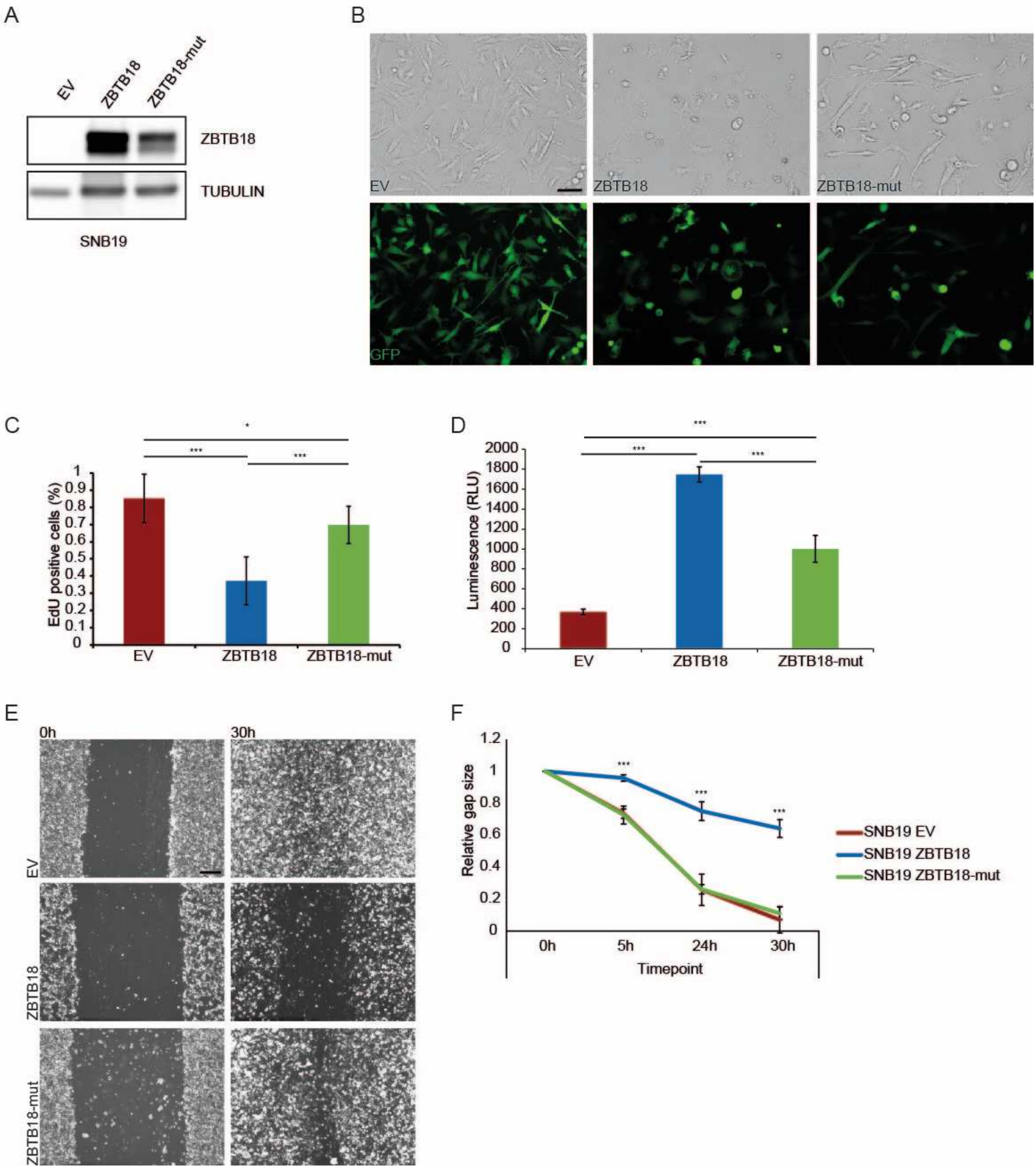

**Figure 1-supplemental figure 2. ZBTB18 and ZBTB18-mut expression differentially alter the phenotype and the properties of the tumour cells.** (**A**) Western blot analysis of FLAG-ZBTB18 expression in SNB19 cells transduced with EV, FLAG-ZBTB18, or FLAG-ZBTB18-mut. (**B**) Micrographs showing the morphology of SNB19 EV controls compared to SNB19 overexpressing ZBTB18 or ZBTB18-mut. Top row is transmitted light, bottom row is GFP; scale bar: 100µm. (**C**) Quantification of the EdU labelling for proliferating cells of SNB19 cells transduced with EV, FLAG-ZBTB18, or FLAG-ZBTB18-mut. n=10; error bars ± s.d. *p < 0.05, **p < 0.01, ***p < 0.001. (**D**) Quantification of the apoptosis by assessing the activity of caspases 3 and 7 in SNB19 cells transduced with EV, FLAG-ZBTB18, or FLAG-ZBTB18-mut. n=4; error bars ± s.d. *p < 0.05, **p < 0.01, ***p < 0.001. (**E**) Micrographs showing the wound closure process in a migration assay with SNB19 cells transduced with EV, FLAG-ZBTB18, or FLAG-ZBTB18-mut. Scale bar: 400µm. (**F**) Quantification of the migration assay described in (E). n=4; error bars ± s.d. *p < 0.05, **p < 0.01, ***p < 0.001.

Figure 2-figure supplement 1

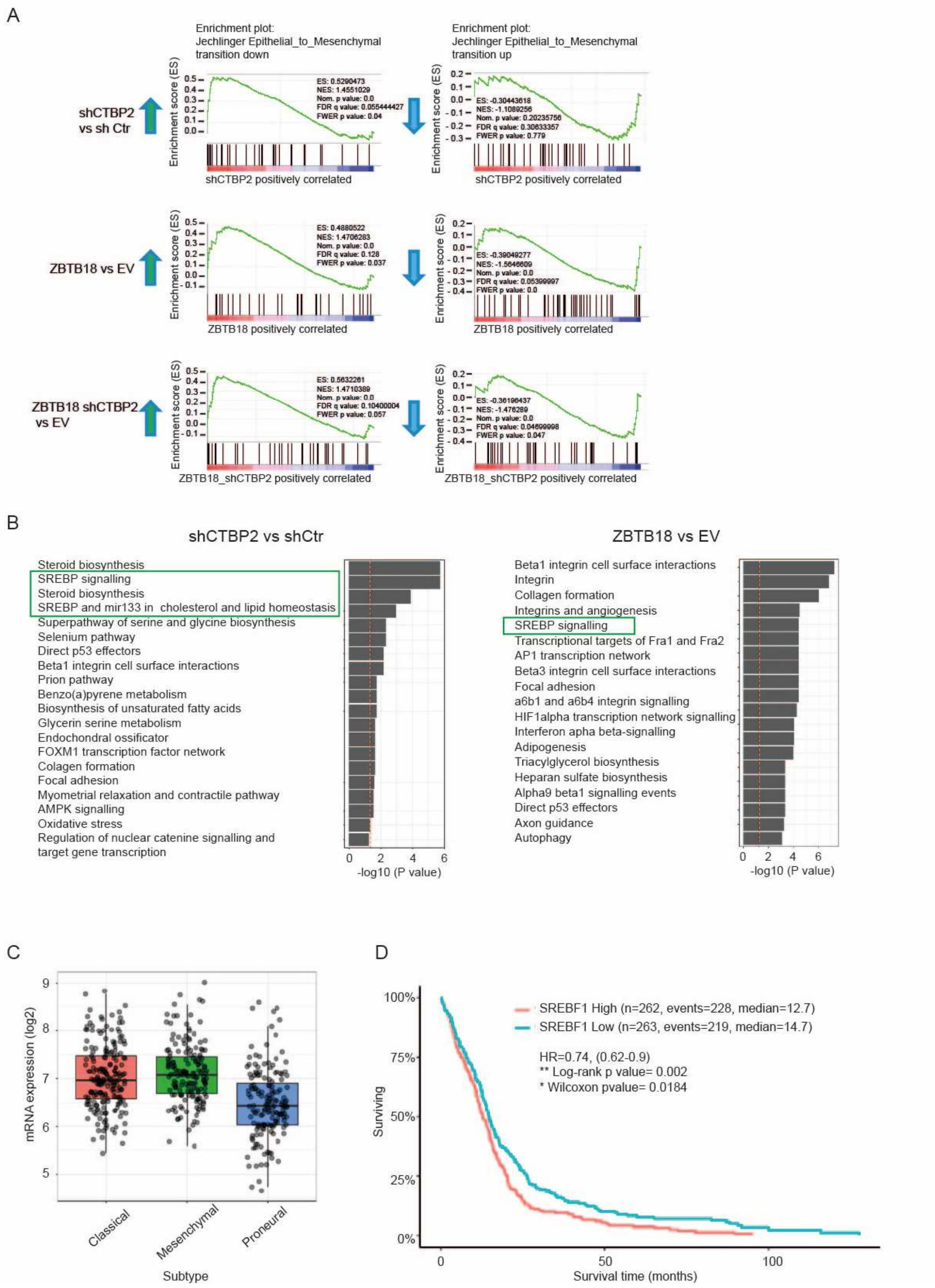

**Figure 2-figure supplement 1. ZBTB18 and CTBP2 regulate EMT and SREBP genes**. (**A**) Gene set enrichment showing upregulation of EMT downregulated genes upon ZBTB18 overexpression and CTBP2 knockdown (left panels) and downregulation of EMT upregulated genes following ZBTB18 overexpression and CTBP2 knockdown (right panels). (**B**) Top 20 downregulated consensus pathways in shCTBP2 vs. shCtr (left panel) and ZBTB18 vs. EV (right panel) from Fisher's exact test comparing the DEGs (adj. pvalue < 0.05, FC < -0.5) to the whole set of quantified genes. Processes related to SREBP signalling are highlighted. (**C**) Box plot showing the expression of SREBF1 in the three GBM subtypes (mesenchymal, classical and proneural) from the TCGA database. The Gliovis platform was used for the analysis. (**D**) Kaplan-Meier plot of TCGA survival data for GBM patients with low versus high SREBF1 expression. The analysis was performed using the Gliovis platform.

Figure 2-figure supplement 2

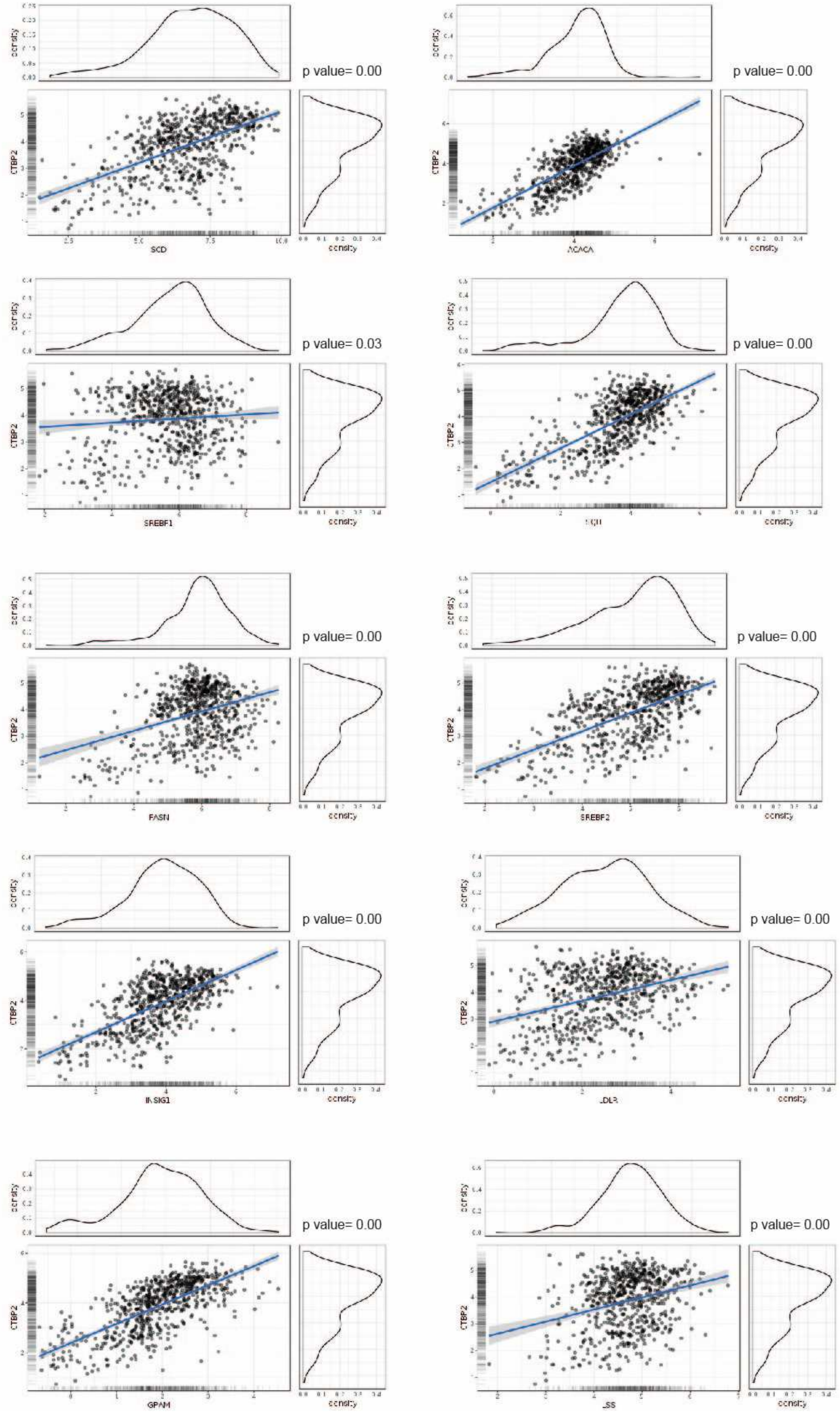

**Figure 2-figure supplement 2. CTBP2 positively correlates with the expression of SREBP genes.** Correlation analysis between CTBP2 and SREBP genes, plotted using the GlioVis portal. RNAseq data from the Chinese Glioma Genome Atlas (CGGA) project (including GBM and low grade gliomas) were used.

Figure 2-figure supplement 3

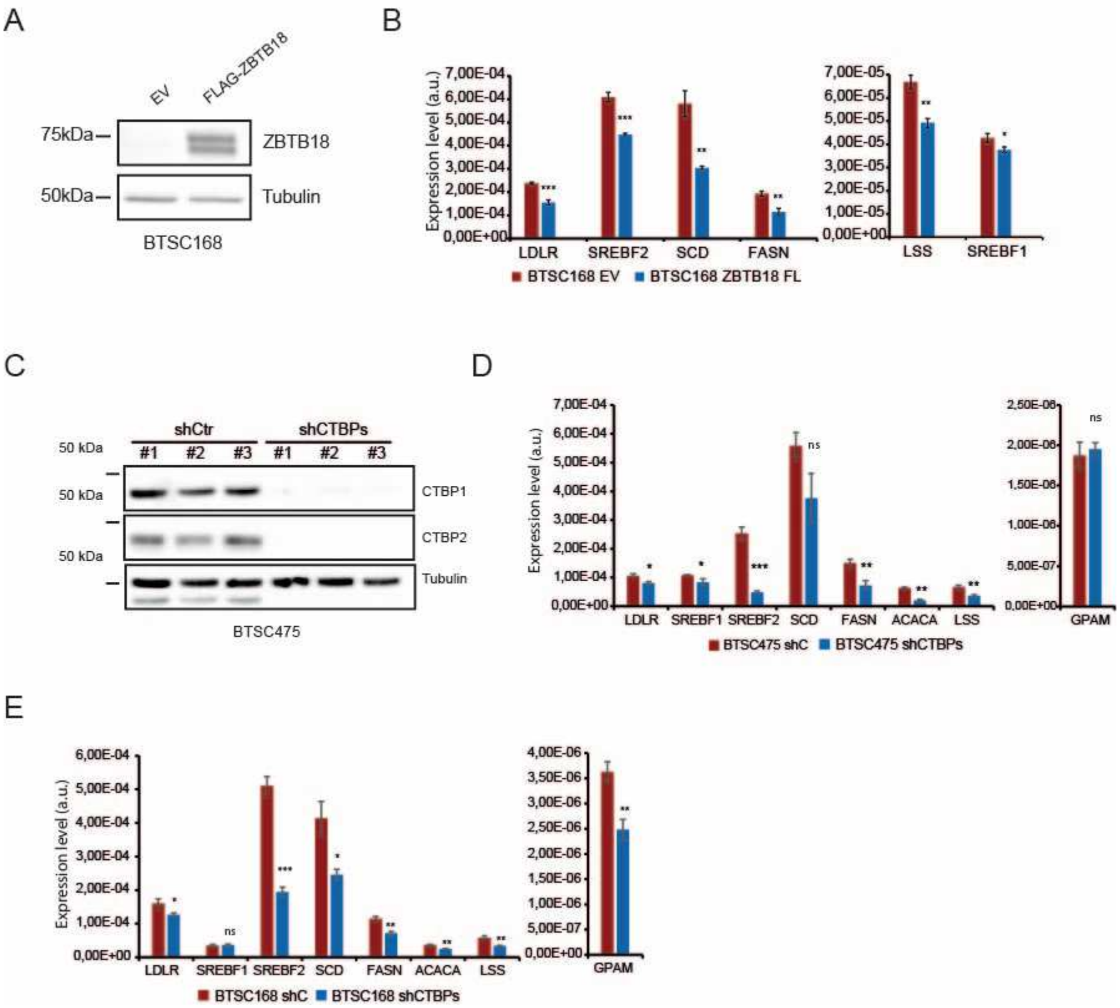

**Figure 2-figure supplement 3. ZBTB18 overexpression and CTBP2 inhibition affect SREBP genes expression**. (**A**) Western blot showing the ectopic expression of ZBTB18 in BTSC168. (**B**) q-RT PCR analysis of SREBP targets in BTSC168 transduced with empty vector (EV) or ZBTB18. Results are presented as the mean of n=3 biological replicates; error bars ± s.d. *p < 0.05, **p < 0.01, ***p < 0.001 by Student's t-test. (**C**) WB analysis of CTBP1 and CTBP2 expression in BTSC475 transduced with shCTBP1 (#8) and shCTBP2 (#1). (**D**) q PCR analysis of selected SREBP genes in BTSC475 cells treated as in (**C**). n=3 biological replicates; error bars ± s.d. *p < 0.05, **p < 0.01, ***p < 0.001 by Student's t-test. (**E**) q-RT PCR analysis of SREBP targets in BTSC168 treated transduced with shCTBP1 (#8) and shCTBP2 (#1) expressing lentivirus. Results are presented as the mean of n=3 biological replicates; error bars ± s.d. *p < 0.05, **p < 0.01, ***p < 0.001 by Student's t-test.

Figure 3-figure supplement 1

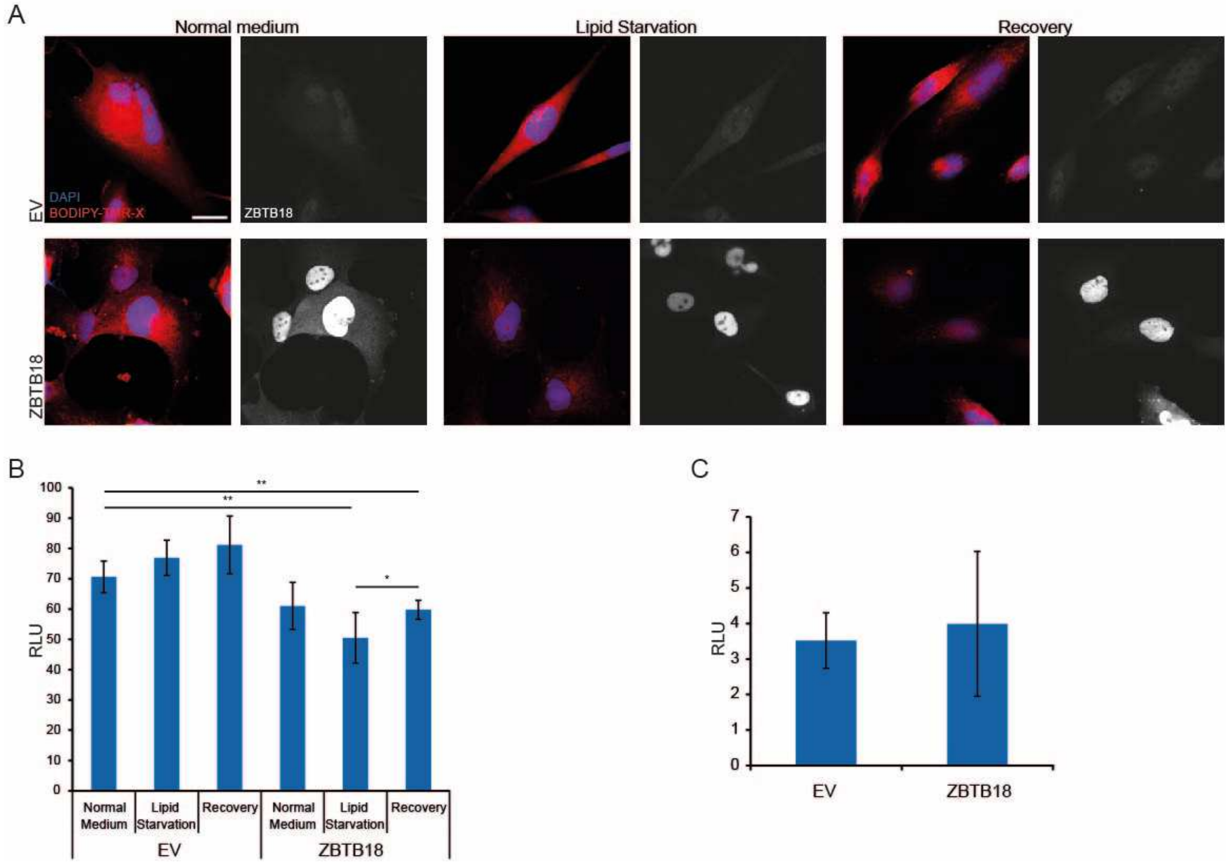

**Figure 3-figure supplement 1. ZBTB18 overexpression affects cell lipid droplets content without affecting lipid uptake.** (**A**) Bodipy TMR-X lipid staining of SNB19 cells expressing empty vector (EV) or ZBTB18, grown in lipid-containing medium (Normal medium), in conditions of lipid starvation (Lipid starvation) or in lipid-containing medium after a period of lipid starvation (Recovery). ZBTB18 was also immunolabeled and nuclei were counterstained with DAPI. Scale bar: 20μm. (**B**) Quantification of the lipid staining shown in (*A*). n=5 biological replicates; error bars ± s.d. $*p < 0.05$, $**p < 0.01$, $***p < 0.001$. (**C**) Quantification of Bodipy-C16 uptake in SNB19 cells expressing EV, ZBTB18 or ZBTB18-mut. n=3 biological replicates; no significance difference was observed.

Figure 4.figure supplement 1

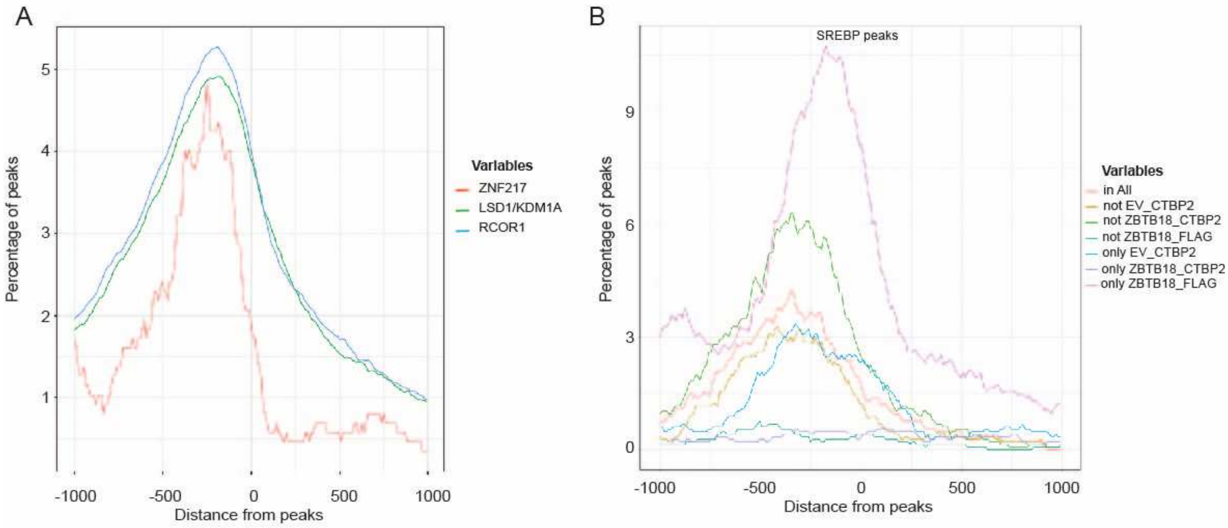


C

| | real | random median | random min | random max | pv | ratio |
|---|---|---|---|---|---|---|
| ZNF217 | 86 | 46 | 28 | 68 | 0 | 1,26470588 |
| KDM1A | 2648 | 1590 | 1474 | 1695 | 0 | 1,56224189 |
| RCOR1 | 2855 | 1856 | 1738 | 1953 | 0 | 1,46185356 |

| | real | random median | random min | random max | pv | ratio |
|---|---|---|---|---|---|---|
| onlyEV_CTBP2 | 115 | 20 | 8 | 34 | 0 | 3,38235294 |
| onlyZNF238_FLAG | 359 | 46 | 24 | 68 | 0 | 5,27941176 |
| onlyZNF238_CTBP2 | 29 | 24 | 9 | 38 | 0,13 | 0,76315789 |
| notEV_CTBP2 | 113 | 9 | 1 | 18 | 0 | 6,27777778 |
| notZNF238_FLAG | 24 | 4 | 0 | 9 | 0 | 2,66666667 |
| notZNF238_CTBP2 | 208 | 16 | 2 | 30 | 0 | 6,93333333 |
| inAll | 113 | 12 | 2 | 22 | 0 | 5,13636364 |

**Figure 4-figure supplement 1. ZBTB18 and CTBP2 peaks overlap with LSD1, ZNF217, NCOR1 and SREBF1 enriched regions from ReMap.** (**A**) ReMap graphics showing enrichment of published ZNF217, LSD1/KDM1A, NCOR1 and CTBP2 peaks at the ZBTB18 and CTBP2 enriched regions. (**B**) ReMap graphics showing enrichment of published SREBF1 peaks at the ZBTB18 and CTBP2 enriched regions. (**C**) Tables showing the number of peaks detected by ReMap analysis in (**A**), top panel and (**B**), bottom panel, compared to random regions. P values are calculate as described in the method sections.

Figure 4-figure supplement 2

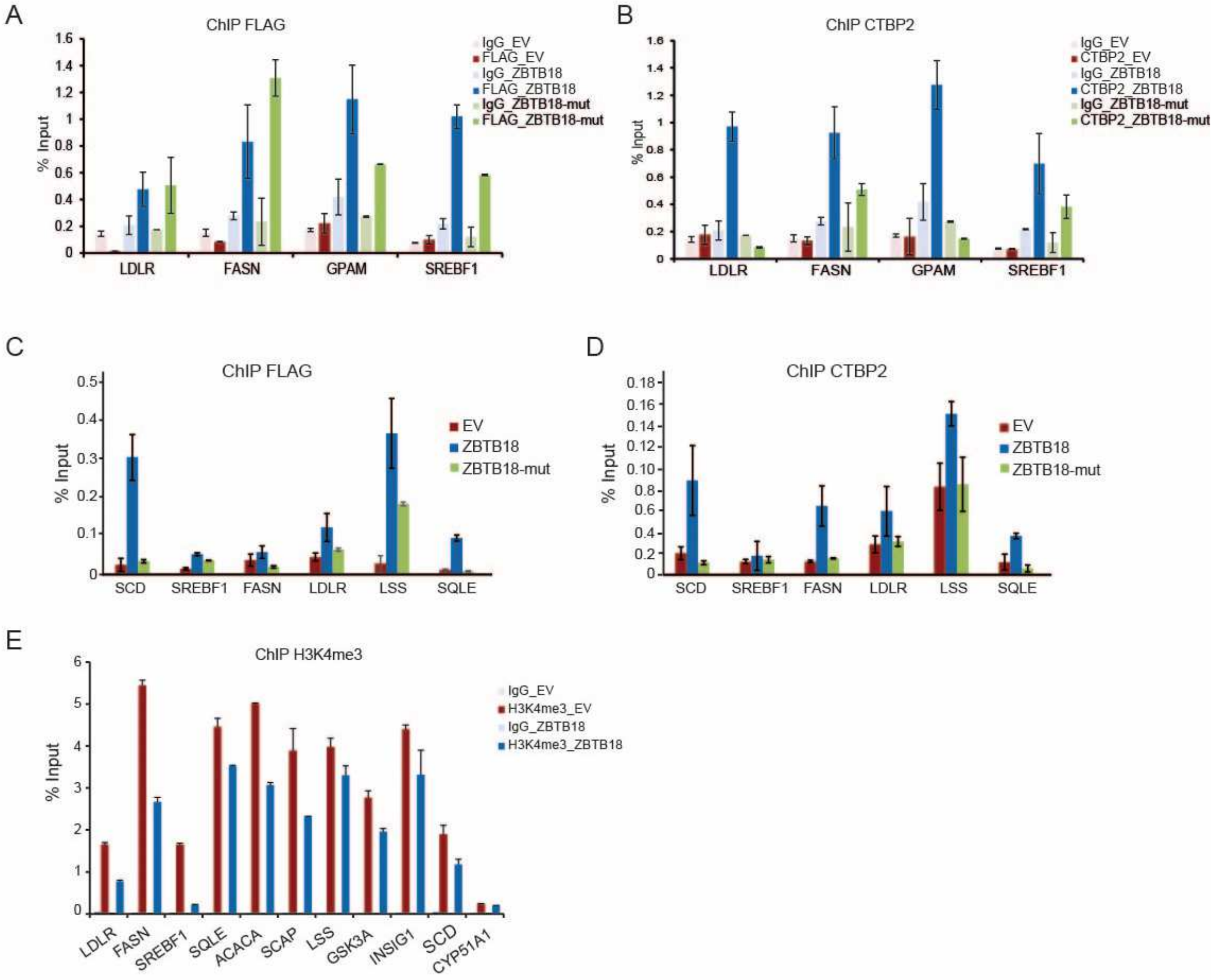

**Figure 4-figure supplement 2. ZBTB18 facilitates the recruitment of CTBP2 at the promoter of SREBP genes. (A-B**) FLAG-ZBTB18 (**A**) and CTBP2 (**B**) ChIP was performed in SNB19 cells, transduced with empty vector (EV), FLAG-ZBTB18 or FLAG-ZBTB18mut, using control beads alone (IgG), anti-FLAG antibodies or anti-CTBP2 antibody. Graphs show representative qPCR results (n=3 technical replicates) of three biological replicates and are expressed in % input as indicated. (**C-D**) Representative qPCR showing binding of FLAG-ZBTB18 (**C**) or CTBP2 (**D**) at selected SREBP gene promoters in BTSC168 cells transduced with empty vector (EV), ZBTB18 and ZBTB18-mut. CTBP2 binding increase in presence of ZBTB18 but not when ZBTB18-mut is expressed. (**E**) H4K4me3 enrichment at the promoter of the indicated SREBP genes in SNB19 cells transduced with empty vector (EV) or FLAG-ZBTB18. Decreased H3K4me3 is consistent with gene repression.

Figure 5.figure supplement 1

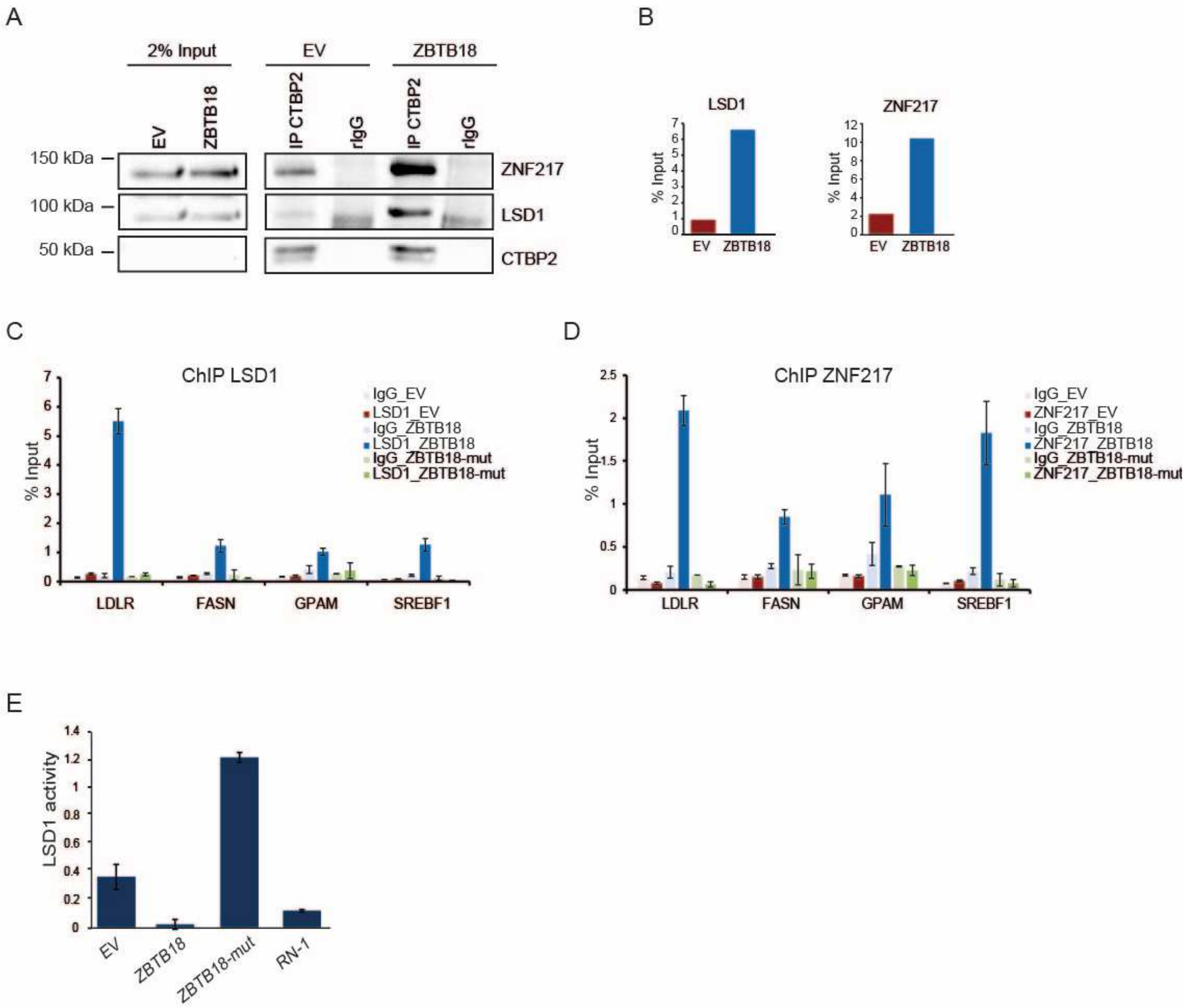

**Figure 5-figure supplement 1. ZBTB18 facilitates the recruitment of LSD1 and ZNF217 at the promoter of SREBP genes.** (**A**) WB analysis of CTBP co-IP in SNB19 cells transduced with EV or FLAG-ZBTB18. (**B**) Quantification of co-IP results. **(C-D**) LSD1 (**C**) and ZNF217 (**D**) ChIP performed using control beads alone (IgG), anti-LSD1 or anti-ZNF217 antibody, in SN19 cells. Graphs show representative qPCR results (n=3 technical replicates) of three biological replicates and are expressed in % input as indicated. (**E**) Bar chart showing LSD1 activity in SNB19 cells transduced with EV, ZBTB18, ZBTB18-mut (all n=7), or treated with the LSD1 inhibitor RN-1 (n=4). LSD1 activity is reduced upon ZBTB18 expression.

#### Supplementary methods

##### Lentiviral Vectors

To produce FLAG-ZBTB18-HisZBTB18, ZBTB18 was PCR amplified from the previously described pCHWMS-eGFP-ZBTB18 lentiviral vector (Fedele et al. 2017) using primers containing a BstXI and PmeI restriction sites. Upon restriction digestion the ZBTB18 fragment was cloned into BstXI and PmeI sites by removing the LUC region of the PCHMWS-eGFP-IRES vector. Primers are listed in the reagent table. ZBTB18 LDL-mut was obtained by site-directed mutagenesis using pCHWMS-eGFP-FLAG-ZBTB18-His as template and the QuikChange II XL Site-Directed Mutagenesis kit (Agilent), according to the manufacturer's instructions. All constructs were sequence validated. Primers are listed in Supplementary Table S2.

Lentiviral stocks production and cell infection were performed as previously described (Fedele et al. 2017). For CTBP1 and CTBP2 knockdown, the following short hairpins RNA cloned in the pLKO lentiviral vector (Sigma Aldrich) were used: shCTBP2-#1(TRCN0000013745), shCTBP2#2 (#TRCN0000013747), shCTBP1#8 (TRCN0000013739) and shCTBP1#9 (TRCN0000273844), or a non-targeting shRNA (SHC002, here shCtr). LSD1 silencing was achieved by lentiviral transfection with MISSION® pLKO.1-puro Empty Vector Plasmid DNA (Sigma Aldrich) harboring either the sequence targeting human LSD1 (TRCN0000046071), or a non-targeting shRNA (SHC002, here shCtr).

For ZBTB18 KO, 4 sgRNA targeting exon 2 (ZBTB18 KO#1 AAAGTCGAGAGTCTCTCCGA; ZBTB18 KO#2 ATCTGCCGAATCCCTCACGG; ZBTB18 KO#3 CAAGCAGGAGAGCGAAAGCG and ZBTB18 KO#4 AAGTGTGAGCACTAATAACA) were cloned in the pKLV-U6gRNA(BbsI)-PGKpuro2BFP

(Addgene, #50946) and lentiviral particles were prepared as described above to transduce BTSC475 cells. Cells were selected with 1µg/mL puromycin, transduced with Cas9-expressing lentiviral particle (pLentiCas9-GFP, Addgene #86145) and selected with 5µg/mL blastacidin. ZBTB18 knockout was verified by Sanger sequencing and Western blot.

LSD1 KO in GBM#22 were generated by the Cogentech Genome Editing Facility (Milan, Italy) and kindly provided by Dr. Pelicci. 2 different sgRNA Target Sequence directed to three different exons were used. LSD1 KO#1:GGTAATTATTATAGGCTCTG (E7T13) (exon 7) and LSD1 KO#3: CTAAATAACTGTGAACTCGG (E6T1)(exon6). Guide RNAs were cloned in all-in-one PX458 lentiviral vector using Nucleofection (kit V, program T-020). GFP positive cells were sorted after 48h and single clones were established by limiting dilution. After single clone propagation, LSD1 knockout was verified by NGS (Ion Proton) and Western blot.

**Quantitative ChIP**

Quantitative ChIP was performed as follows: SNB19 or BT168 cells were seeded at $1x10^6$ cells confluency and transduced either with the control or ZBTB18 expressing lentiviral vectors as described before. After 72 hours cells were washed in PBS and incubated in the presence of ChIP Crosslink Gold solution (Diagenode) for 30 minutes at room temperature, fixed by addition of 1% formaldehyde for 20 minutes at room temperature and quenched with PBS for 5 minutes. The cells were resuspended in 2 ml L1 buffer (50 mM Tris pH 8.0; 2 mM EDTA pH 8.0; 0,1% NP40; 10% glycerol; protease inhibitors) per $10^7$ cells, and lysed on ice for 5-10 minutes. The nuclei were collected by centrifugation at 800g for 5 minutes at 4ºC and lysed in 1 ml of L2 buffer (0,2% SDS; 10 mM EDTA; 50 mM Tris pH 8; protease inhibitors). The suspension was sonicated in a cooled Bioruptor

Pico (Diagenode), and cleared by centrifugation for 10 minutes at 13000 rpm. The chromatin (DNA) concentration was quantified using NanoDrop (Thermo Scientific) and the sonication efficiency monitored on an agarose gel. Protein A sepharose (PAS) beads (GE Healthcare) were blocked with sonicated salmon sperm DNA (200 mg/ml beads) and BSA (250 mg/ml beads) in dilution buffer (0.5% NP40; 200 mM NaCl; 50 mM Tris pH 8.0; protease inhibitors) for 2 hours at room temperature. The chromatin was diluted 10x in the dilution buffer and pre-cleared with blocked PAS for 1 hour at 4°C. 5 ug of pre-cleared chromatin was incubated with 5 ug of antibody O/N at 4°C, then with 40 ul of blocked PAS further 2 hours at 4˚C. Antibodies are listed in Supplementray Table S7. Control mock immunoprecipitation was performed with blocked PAS. The beads were washed 4x with WB-250 (0.02% SDS; 0.5% NP40; 2 mM EDTA; 250 mM NaCl; 20 mM Tris pH 8.0). The immuno-complexes were eluted by two 15 minutes incubations at 30°C with 100 µl elution buffer (1%SDS, 100mM $NaHCO_3$), and de-crosslinked overnight at 65°C in the presence of 10U RNase (Roche). The immunoprecipitated DNA was then purified with the QIAquick PCR purification kit (Qiagen) according to manufacturer's protocol and analyzed by qRT-PCR. Primers are listed in SupplementaryTable S4.

**MS analysis**

For MS analysis of BTSC268, one ug of peptides were analyzed on a Q-Exactive Plus mass spectrometer (Thermo Scientific, San Jose, CA) coupled to an EASY-nLCTM 1000 UHPLC system (Thermo Scientific). The analytical column was self-packed with silica beads coated with C18 (Reprosil Pur C18-AQ, d = 3 Â) (Dr. Maisch HPLC GmbH, Ammerbusch, Germany). For peptide separation, a linear gradient of increasing buffer B (0.1% formic acid in 80% acetonitrile, Fluka) was applied, ranging from 5 to 40% buffer B over the first 90 min and from 40 to 100% buffer B in the subsequent 30 min (120 min separating gradient length). Peptides were analyzed in data dependent acquisition mode (DDA). Survey scans were performed at 70,000 resolution, an AGC target of 3e6 and a maximum injection time of 50 ms followed by targeting the top 10 precursor ions for fragmentation scans at 17,500 resolution with 1.6 m/z isolation windows, an NCE of 30 and an dynamic exclusion time of 35 s. For all MS2 scans the intensity threshold was set to 1.3e5, the AGC to 1e4 and the maximum injection time to 80 ms. Raw data were analyzed with MaxQuant (v 1.6.14.0) allowing two missed cleavage sites, no variable modifications, carbamidomethylation of cysteines as fixed modification, using label free quantification (LFQ), and match between runs (MBR) set to activated. The Human-EBI-reference database was downloaded from https://www.ebi.ac.uk/ on Jan 9th 2020. Only unique peptides were used for quantification.

**SupplementaryTable 2.** List of primers used for cloning and site-directed mutagenesis.

| **Primer name** | **Primer sequence** |
|---|---|
| ZNF238_LDLmut_f | CTGAAAGGCTGGACTTGACAGACGAGGCCGACACACAGTCAACATCTGCCGAAT |
| ZNF238_LDLmut_r | ATTCGGCAGATGTTGACTGTGTGTCGGCCTCGTCTGTCAAGTCCAGCCTTTCAG |
| BstXI-flag-hZBTB18-sense | TGGCCACAACCATGGACTACAAGGACGACGATGACAAGTGTCCTAAAGGTTATGAAGACAG |
| PmeI-HA-hZBTB18-antisense | GCCTTGGTTTAAACTTAAGCGTAATCTGGAACATCGTATGGGTATTTCCAAAGTTCTTGAGAGCTA |

**SupplementaryTable 3.** List of primers used for qRT-PCR.

| **Primer name** | **Primer sequence** |
|---|---|
| 18s_F | CGCCGCTAGAGGTGAAATTC |
| 18s_R | CTTTCGCTCTGGTCCGTCTT |
| ABCA1_F1 | ACCCACCCTATGAACAACATGA |
| ABCA1_R1 | GAGTCGGGTAACGGAAACAGG |
| LDLR_F1 | TCTGCAACATGGCTAGAGACT |
| LDLR_R1 | TCCAAGCATTCGTTGGTCCC |
| SREBF1_F1 | ACAGTGACTTCCCTGGCCTAT |
| SREBF1_R1 | GCATGGACGGGTACATCTTCAA |
| INSIG1_F2 | ATCCAGAGGAATGTCACTCTCTT |
| INSIG1_R2 | AGGGGTACAGTAGGCCAACAA |
| ACACA_F1 | ATGTCTGGCTTGCACCTAGTA |
| ACACA_R1 | CCCCAAAGCGAGTAACAAATTCT |
| FASN_F2 | CCGAGACACTCGTGGGCTA |
| FASN_R2 | CTTCAGCAGGACATTGATGCC |
| SCD_F2 | GCCCCTCTACTTGGAAGACGA |
| SCD_R2 | AAGTGATCCCATACAGGGCTC |

**SupplementaryTable 4.** List of primers used for quantitative ChIP.

| **Primer name** | **Primer sequence** |
|---|---|
| LDLR_ChIP_TSS_Fw1 | ATAGAAAGTGGCGGAAGTTCC |
| LDLR_ChIP_TSS_Rv1 | ATAGAAAGTGGCGGAAGTTCC |

| | |
|---|---|
| FASN_ChIP_Fw | GGACGAAATGGGGATAGCCTA |
| FASN_ChIP_Rv | CTGTGGTGTGTGGGTTGGTAT |
| GSK3A_Fw | GGAAAGGCATCTGTCGGGG |
| GSK3A_Rv | GAGTGGCTACGACTGTGGTC |
| INSIG1_ChIP_fw | CCTTCCTCGCTCTTTGTCTCT |
| INSIG1_ChIP_rv | GTTGATCATCTCCCCAACCTT |
| SCAP_ChIP_fw | TGAGGTCATAAACCCACTCAGA |
| SCAP_ChIP_rv | TAGAACCTGCTTTGGTGCTGT |
| LSS_ChIP_fw | TGCGTGGTTTAGAGATGAAGG |
| LSS_ChIP_rv | CTGGACACCGTAAGTTGCTTC |
| ACACA_ChIP_fw | AGTTCCCTCAGCCTCAATTTC |
| ACACA_ChIP_rv | CTGACTTTTGATCCGACCAGT |
| CYP51A1_ChIP_fw | TGAGTCTTTGGCTTTCGTACC |
| CYP51A1_ChIP_rv | CAAGACAATCCCACCAAGATG |
| SQLE-Z-fw | TGCGACGGTTACTCTGGTTAC |
| SQLE-Z-rv | CCAGGGTACCTCCCTCAGAT |
| SREBF1-fw1 | CCCTCTGTAATGGTGTGCCTA |
| SREBF1-rv1 | AAGCGCTCAGCAAGTAAACTG |

**SupplementaryTable 5.** List of antibodies used for western blots.

| **Antibody name** | **Company** |
|---|---|
| mouse anti-FLAG | Sigma #F1804 |
| rabbit anti FLAG | (Cell Signaling #2368S) |
| rabbit anti-ZBTB18 | AbCam #ab118471 |
| mouse anti-CTBP2 | BD Biosciences #612044 |
| mouse anti-CTBP1 | BD Biosciences #612042 |

| | |
|---|---|
| rabbit anti-CTBP2 | Cell Signaling #13256S |
| rabbit anti-CTBP1 | Cell Signaling #8684 |
| rabbit anti-LSD1 | Millipore #17-10531 |
| mouse anti-LSD1 | Santa Cruz # sc-53875 |
| rabbit anti-ZNF217 | Thermo Fisher scientific #720352 |
| mouse anti alpha tubulin | Abcam #ab7291 |

**SupplementaryTable S6.** List of antibodies used for co-IP.

| **Antibody name** | **Company** |
|---|---|
| rabbit anti-CTBP2 | Cell Signaling #13256S |
| mouse anti-CTBP | Santa Cruz #sc-17759 |
| mouse anti-LSD1 | Santa Cruz # sc-53875 |
| mouse anti-FLAG | Sigma # F1804 |
| rabbit anti-ZBTB18 | AbCam #ab118471 |
| rabbit anti-ZBTB18 | Proteintech #12714-1-AP |
| normal mouse IgG | Santa Cruz, #sc-2025 |
| normal rabbit IgG | Santa Cruz, #sc-2027 |

**SupplementaryTable S7.** List of antibodies used for ChIP.

| **Antibody name** | **Company** |
|---|---|
| rabbit anti-CTBP2 | Active Motif #61262 |
| mouse anti-FLAG | Sigma #F1804 |
| rabbit anti-LSD1 | Merck Millipore #17-10531 |
| rabbit anti-ZNF217 | Thermo Fisher Scientific #720352 |
| rabbit anti-H3K4me3 | Diagenode #C15410003 |
| rabbit anti-H3K4me2 | Cell Signaling #9725 |
| rabbit anti-H3K9me2 | Cell Signaling #9753 |
| mouse anti-H3K9me2 | Abcam #ab1220 |
